# Rapid generation of ventral A9-like dopaminergic neurons from patterned iPSCs

**DOI:** 10.1101/2025.10.31.685897

**Authors:** Kriti Chaplot, Lin Zhang, Julia Wessman, Miguel Rivera, Zhihui Wang, Fei Wang, Xin Duan, Pamela M. England, Iain C. Clark, Erik M. Ullian

## Abstract

*In vitro* modelling of highly vulnerable nigral dopaminergic (DA) neuronal subtypes in Parkinson’s disease (PD), is necessary for studying disease mechanisms. Here, we optimized a new approach by expressing the pioneer neurogenic transcription factor, *Achaete-scute-like 1* (*Ascl1*), implicated in determining dopaminergic fate. Sequential small-molecule patterning of iPSCs into early floor plate mesencephalic progenitors, followed by inducible *Ascl1* expression, rapidly differentiates midbrain DA neurons. Immunocytochemistry and transcriptomic analysis of these patterned Ascl1-driven DA neurons (PA-DANs) confirmed midbrain-lineage specificity. Importantly, we found an enrichment of DA subpopulations that corresponded to the adult human ventral SOX6-positive A9 DA subtypes vulnerable in PD. Furthermore, we combined these ventral A9-like PA-DANs with human iPSC-derived midbrain astrocytes and microglia in defined ratios to generate mature 3D A9-like assembled organoids that display characteristic spontaneous neuronal activity and electrical propagation along the axon. Our method efficiently generates a mature and functional A9-like DA neuronal platform to study PD.

**Highlights:**

- Sequential midbrain patterning and Ascl1 expression accelerates DA differentiation
- PA-DANs resemble human adult ventral A9-like DA subtypes vulnerable in PD
- 3D assembled organoids show mature identity of PA-DANs, iAstrocytes and iMicroglia
- PA-DANs matured in 3D organoids show neuronal network activity within weeks

**eTOC blurb:** In this study, Ullian and colleagues have developed a rapid method to differentiate dopaminergic neurons, using small molecules to generate early floor plate mesencephalic progenitors from human iPSCs and sequentially expressing a pioneer transcription factor, Ascl1, that accelerates uniform dopaminergic neurogenesis. Patterned Ascl1-driven dopaminergic neurons (PA-DANs) in 2D and 3D assembled organoids serve as a platform to study Parkinson’s disease

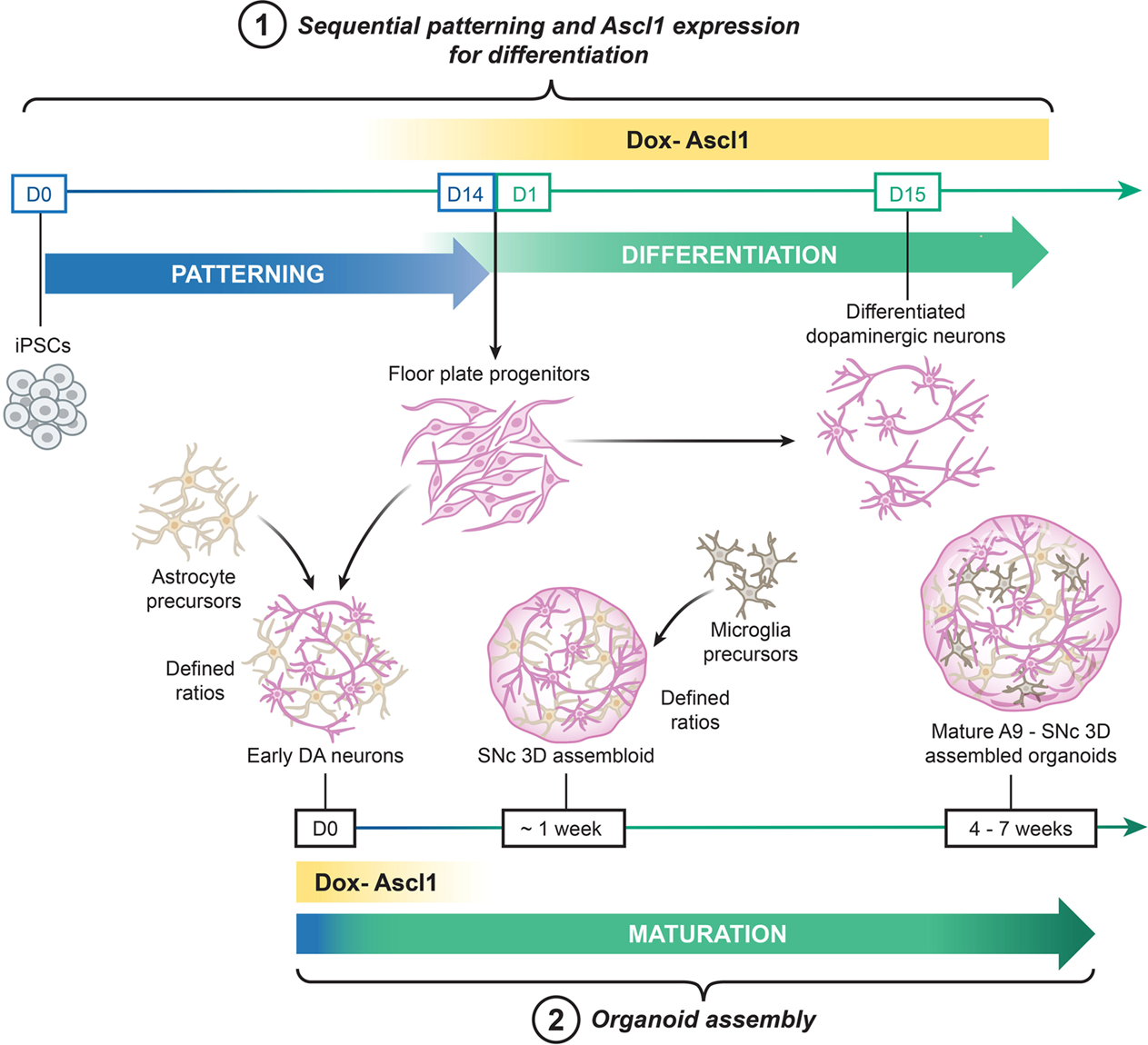

## Introduction

Dopaminergic (DA) subregional heterogeneity is consequential to disease vulnerability and selective cell death. DA neurons of the human adult substantia nigra pars compacta (SNc), corresponding to the ventral A9 region, are marked by the expression of *SOX6* and *AGTR1*, express higher levels of PD-causative genes and have been shown to be more vulnerable to Parkinson’s disease (PD) (Kamath et al., 2022). On the other hand, both the dorsal A9 region of the SNc and the neighboring ventral tegmental area (A10), contain DA neurons that highly express *CALB1* and are more resilient to neuronal death in PD (Kamath et al., 2022). Recapitulating DA subpopulations specific to the SNc, enriched in vulnerable DA subregional subtypes, using *in vitro* human cell-based systems is vital in studying human neurodegenerative disease mechanisms and the causes of selective cell vulnerability.

Human induced pluripotent cells have been successfully used to derive different cell types of the brain to mimic and study human neuron-neuron and neuron-glia networks. In the context of studying iPSC-derived DA neurons, there have been two major approaches to develop dopaminergic neurons from iPSCs – small-molecule-mediated directed differentiation and inducible transcription factor-driven differentiation (Kim et al., 2021; Kriks et al., 2011; Powell et al., 2021; Sarrafha et al., 2021; Theka et al., 2013).

Using the directed differentiation approach, *in vitro*, midbrain specificity has been achieved by exposing iPSCs to additional small molecules, growth and neurotrophic factors that regulate key developmental pathways such as sonic hedgehog (SHH), fibroblast growth factor 8 (FGF8) and Wingless-related integration site (Wnt) signaling, along with dual SMAD inhibition of Bone Morphogenetic Protein (BMP) and Tumor Growth Factor β (TGFβ) pathways, to pattern floor plate mesencephalic progenitors that further differentiate into DA neurons with prolonged use of midbrain-specific neurogenic factors (Calatayud and Muñoz-Pedrazo, 2022; Kim et al., 2021; Kriks et al., 2011; Sarrafha et al., 2021). However, this process in both 2D and 3D organoid cultures is slow, time-consuming (>30-60 days) and often yields variable subregional DA heterogeneity, limiting its precision in modeling specific DA subtypes.

In contrast, TF-driven differentiation uses the inducible expression of neuronal transcription factors in iPSCs or embryonic stem cells (ePSCs) to rapidly generate neurons (<10-20 days). For example, NGN2 expression under a doxycycline-inducible promoter primarily yields vGlut1- or vGlut2-positive cortical or motor neurons (Limone et al., 2023; Nehme et al., 2018; Zhang et al., 2013). Ascl1 expression, on the other hand, has been employed to generate GABAergic-like neurons and tyrosine hydroxylase-positive DA neurons (Aydin et al., 2019; Lin et al., 2025; Lundie-Brown et al., 2025; McDonald et al., 2024; Ng et al., 2021; Powell et al., 2021; Vainorius et al., 2023). Ascl1 expression in combination with midbrain-specific transcription factors such as LMX1a/b, FOXA2, Nurr1, Pitx3 and En1, has been used to rapidly produce DA neurons of CNS identity (Ng et al., 2021). However, overall DA neurogenesis and dopaminergic subregional specificity using inducible TF expression is lower than that produced using the directed differentiation approach. Rapid induced differentiation of DA neurons driven by two or more TFs in conjunction with Ascl1, does not take into consideration the vital timing and tight hierarchical TF regulation necessary for subregional subtype development (Ng et al., 2021; Powell et al., 2021; Theka et al., 2013).

A promising yet underexplored strategy to enhance DA subregional specificity is to combine **temporal control** of inducible TF expression with **directed differentiation** by optimizing the timing of TF expression akin to that seen during embryonic development. Given the strong role of Ascl1 in dopaminergic fate determination and its prior utilization in generating dopaminergic neurons, we asked whether inducible Ascl1 expression during midbrain development could be leveraged for rapid and efficient generation of nigral DA subtypes.

Here, we report a new strategy that converges existing protocols of midbrain specification and Ascl1 expression to rapidly and uniformly generate high-quality and high-percentage mature A9-like dopaminergic neurons (∼15-30 days). In our method, we take advantage of the patterning process based on the fundamentals of midbrain development by supplementing the medium with midbrain-specific effectors modulating developmental pathways such as sonic hedgehog, Wnt, FGF8 signaling, in addition to dual SMAD inhibitors, for a short duration of time, to generate early floor plate progenitor cells. We then induce the expression of Ascl1, a transcription factor essential for midbrain neurogenesis, which accelerates the transition and differentiation of these progenitors to neuronal fate. Using single cell RNA-sequencing, we found that Ascl1 expression in patterned midbrain progenitors yields a A9-specific DA neuronal population with genetic signatures comparable to the ventral human adult SNc transcriptome. Finally, we report a unique method to generate 3D midbrain assembled organoids, by combining iPSC-derived midbrain-astrocytes with our patterned *Ascl1*-driven DA neurons (PA-DANs), and successfully integrating iPSC-derived microglia, supporting active neuronal networks and neuron-glia connections that more closely mimic the human SNc.

## Results

### Inducible *Ascl1* expression in iPSC-derived early floor plate progenitors is sufficient to enhance midbrain fate and rapidly differentiate TH-positive DA neurons

We designed a more efficient protocol to develop dopaminergic neurons from iPSCs by focusing on the specificity and maturation of floor plate progenitor population. To do this, we engineered piggybac-human iPSC lines to express *Ascl1*, the neurogenic transcription factor and chromatin modulator, crucial for dopaminergic determination (Earley et al., 2021; Kim et al., 2007), under a doxycycline (DOX)-inducible promoter (Ng et al., 2021) (Figure 1A).

**Figure 1:**
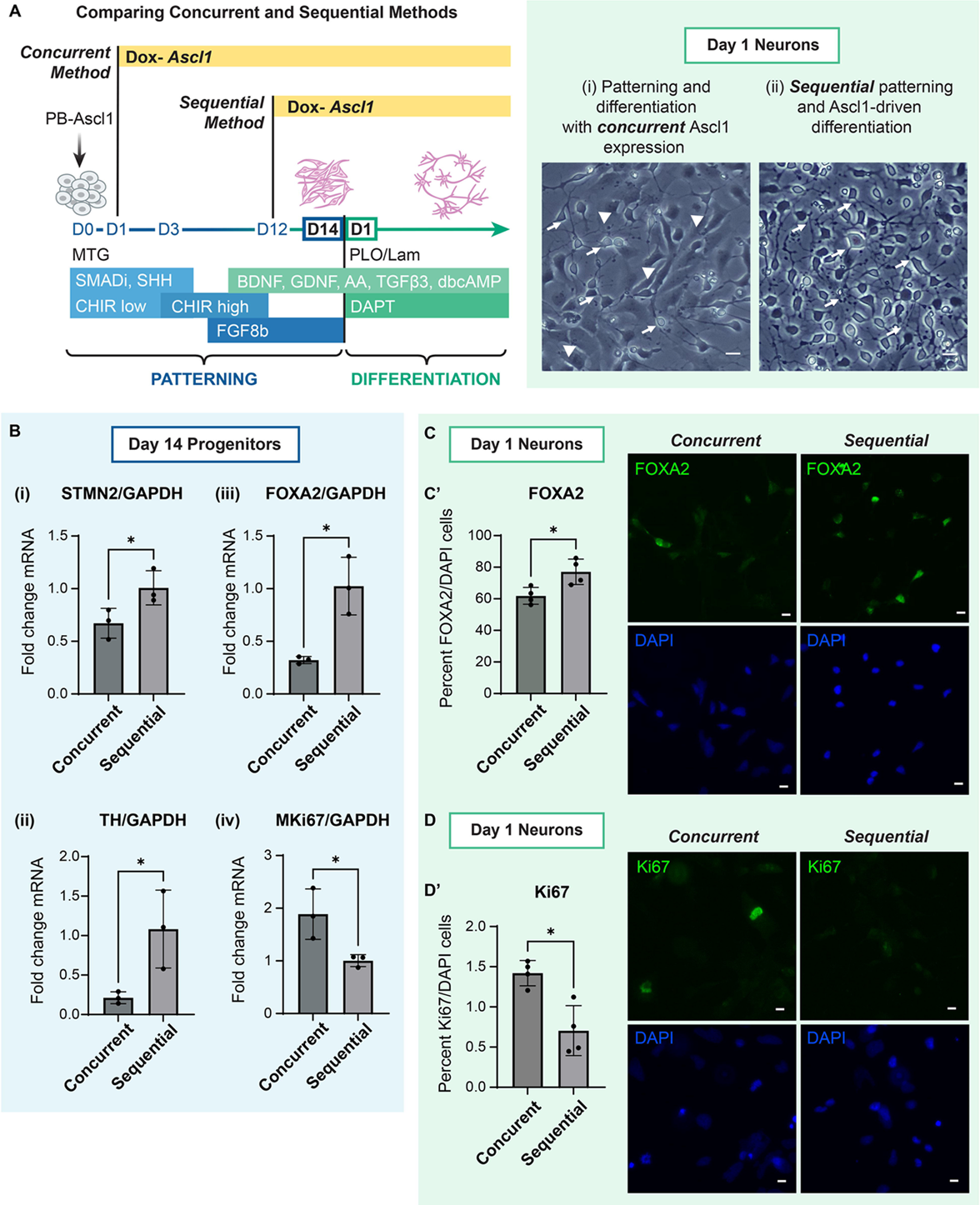
Sequential Ascl1 expression in human iPSC-derived early floor plate mesencephalic progenitors enhances dopaminergic neurogenesis. See also Figure S1. A. Schematic representing midbrain specification of iPSCs into floor plate mesencephalic progenitors which differentiate into dopaminergic neurons, and timelines (left) describing 2 novel approaches to generate DA neurons: (i) concurrent expression of Ascl1 during directed differentiation, (ii) and sequential expression of Ascl1 after exposure to patterning small molecules during directed differentiation. Representative brightfield images (right) of cells generated via the 2 approaches after 24 hours of plating (i) concurrent patterning and Ascl1 induction yielding mixed progenitor-like and neuron-like morphologies, and (ii) sequential patterning cues and Ascl1 induction yielding high neurogenesis. Arrows indicate neuron-like morphology, arrowheads indicate progenitor-like morphology. Scale bar 20µm. B. Graphs showing fold change of normalized mRNA levels of STMN2 (i), TH (ii), FOXA2 (iii) significantly higher and MKi67 (iv) significantly lower in Day 14 progenitors generated via sequential method than via the concurrent method. Welch’s t-test, one tailed. P-value *: <0.05. Data are represented as mean ± SD. N=3. C. Representative images of Day 1 cells generated via concurrent and sequential methods immunostained with midbrain markers, FOXA2 (green) and DAPI (blue) after 24 hours of plating. Scale bar 10µm. Graph (C’) showing significantly high expression of FOXA2 in generated using the sequential method 24 hours after plating. Welch’s t-test, two tailed. P-value *: <0.05. Data are represented as mean ± SD. n=4, 3 fields from each replicate. Approx 2000 total cells counted per condition. D. Representative images of cells generated via concurrent and sequential methods immunostained with cell cycle marker, Ki67 (magenta) and DAPI (blue), after 24 hours of plating. Scale bar 10µm. Graph (D’) showing significantly high expression of Ki67 in cells generated using the sequential method 24 hours after plating. Welch’s t-test, two tailed. P-value *: <0.05. Data are represented as mean ± SD. n=4, 3 fields from each replicate. Approx 2000 total cells counted per condition.

We first tested the neurogenic activity of *Ascl1* in hiPSCs, by using DOX to generate 3-days old *Ascl1*-induced neuronal precursors (Figure S1A, panel i). After 24 hours of plating these precursor cells in midbrain differentiation medium containing DOX, we found limited neurogenesis and a proliferative precursor population, (Figure S1A, panel i). Addition of anti-mitotic agent, AraC, and Notch-inhibitor, DAPT, during the generation of precursors was sufficient to reduce this proliferative population and support Ascl1-induced neurogenesis, as previously reported (Chanda et al., 2014). These data indicated that *Ascl1* expression alone is not sufficient to induce robust neuronal differentiation in hiPSCs as substantial progenitor populations persist.

Having established that Ascl1-induced expression alone is not sufficient to induce robust DA neuron differentiation, we next compared the *Ascl1*-induced neuronal induction to established protocols for directed differentiation of DA neurons. To do this, we patterned hiPSCs using SMAD inhibitors, SHH, Wnt and FGF8b signaling modulators for 14 days to generate early floor plate mesencephalic progenitors (Calatayud and Muñoz-Pedrazo, 2022) (Figure S1A, panel ii). Upon 24 hours of plating the early progenitors in midbrain differentiation medium, the cells remained in the progenitor stage with no discernable neurogenesis (Figure S1A, panel ii). This showed that at 14 days, early progenitors generated using patterning cues were not mature enough for rapid neuronal differentiation.

We next asked if combining these two approaches could achieve both robust differentiation to midbrain lineage and rapid differentiation into DA neurons. We hypothesized that employing inducible Ascl1 during the generation of early floor plate mesencephalic progenitors would enhance progenitor maturation, increase neuronal differentiation and reduce non-neuronal proliferation as is observed in vivo. We did this in two ways, 1) - Concurrent *Ascl1* expression during patterning for 14 days (Figure 1A, panel i) or 2) - Sequential patterning for 12 days followed by *Ascl1* expression for 48 hours (Figure 1A, panel ii). In the concurrent *Ascl1* expression method, 24 hours after plating the Day 14 progenitors in midbrain differentiation medium with DOX, we found increased neurogenesis, however we still identified numerous cells with progenitor-like morphology in this population (Figure 1A, panel i).

Because *Ascl1* expression is known to be upregulated in progenitor cells just prior to DA neuron generation during midbrain development (Kim et al., 2008, 2007), we hypothesized that the induction of *Ascl1*, in pre-patterned 12-day old early progenitors may be sufficient to mimic developmental signals, yielding higher proportions of DA neurons. After 24 hours of differentiating the Day 14 progenitors generated via this sequential *Ascl1* expression method, we found a clear enhancement in neuronal morphology (Figure 1A, panel ii). Thus, we found that, unlike in iPSCs, inducing *Ascl1* expression in early floor plate mesencephalic progenitors was sufficient to trigger their rapid transition into differentiated neurons.

To further compare the efficiency of DA neurogenesis from progenitors generated via the sequential method against the concurrent method, we assessed the expression of neuronal and progenitor markers via quantitative PCR. We found significantly higher mRNA expression of neuronal marker, *STMN2* and DA-specific marker, *Tyrosine hydroxylase* (*TH*) in Day 14 progenitors generated via the sequential method as compared to the concurrent method, indicating Ascl1-driven transition of floor plate progenitors into differentiated neurons (Figure 1B, panel i, panel ii). Additionally, we also found higher midbrain progenitor marker, *FOXA2* mRNA and reduced *MKi67* mRNA in the sequentially-generated progenitors (Figure 1B, panel iii, panel iv). Upon immunostaining, we confirmed that the percentage of cells expressing FOXA2 were significantly higher in cells differentiated generated using the sequential method (88%) than using the concurrent method (62%), while Ki67 was found to be expressed in less than 1% of the population of cells produced via the sequential method, and significantly lower than that produced using the concurrent method (>1%) (Figure 1C, 1C’, 1D, 1D’). These results suggested that sequential induced expression of *Ascl1*, but not concurrent *Ascl1* expression, in early floor plate mesencephalic progenitors, was able to improve midbrain specificity and terminal DA differentiation.

We further tested the efficiency of this method by implementing it across three iPSC lines which we independently engineered with piggybac-Ascl1 to generate PA-DANs (Figure 2A). We found a comparable high (>70%) percentage of cells expressing En1, LMX1 and Nurr1 after 24 hours of plating the patterned progenitors generated from each piggybac-Ascl1 transfected iPSC lines (Figure 2B, panel i, panel ii, panel iii). We also tested the extent of DA neurogenesis 24 hours after differentiating PA-progenitors from different genetic backgrounds and found similarly high percentages of TH-positive DA neurons (Figure 3B, 3B’). This confirmed that our sequential method worked consistently for different iPSC backgrounds and could be easily adapted for inducible Ascl1-expressing stable iPSC lines.

**Figure 2:**
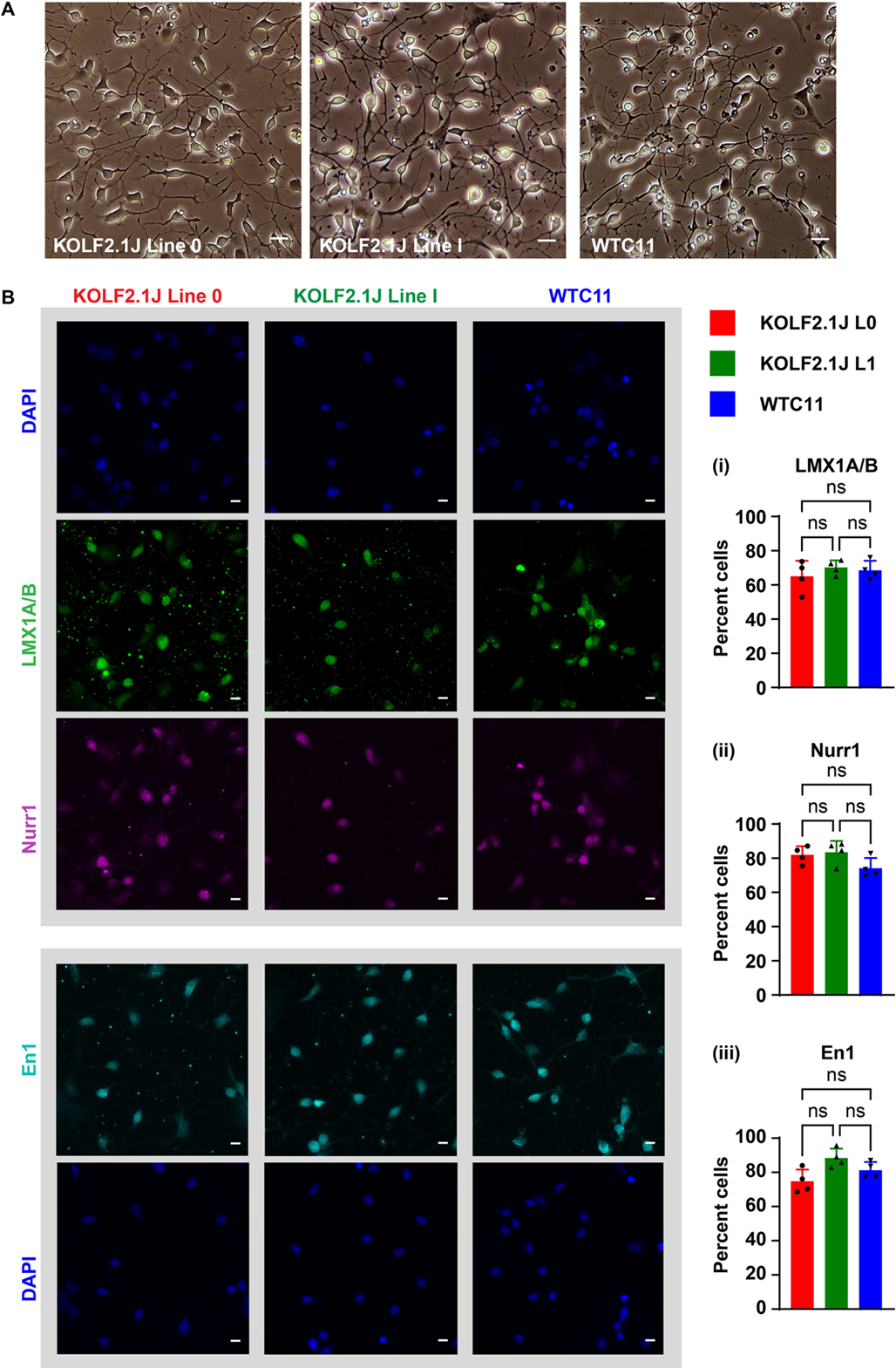
Sequential method to generate PA-DANs can be implemented consistently in different iPSC lines. A. Representative brightfield images of PA-DANs generated from KOLF2.1J lines (Line 0, Line 1) and WTC11 iPSCs independently engineered with piggybac-Ascl1. Scale bar 20µm. B. Representative images of PA-DANs generated from different iPSC lines immunostained with LMX1A/B (green), Nurr1 (magenta) and En1 (cyan) with corresponding images of DAPI (blue) nuclear staining. Scale bar 10µm. Graphs showing percentage of PA-DANs expressing LMX1A/B (i), Nurr1 (ii) and En1 (iii) in different iPSC lines as indicated. One-way ANOVA, multiple comparison, two tailed, ns: not significant. Data are represented as mean ± SD. n=4, 3 fields from each replicate. Approx 2000 total cells counted per condition.

**Figure 3:**
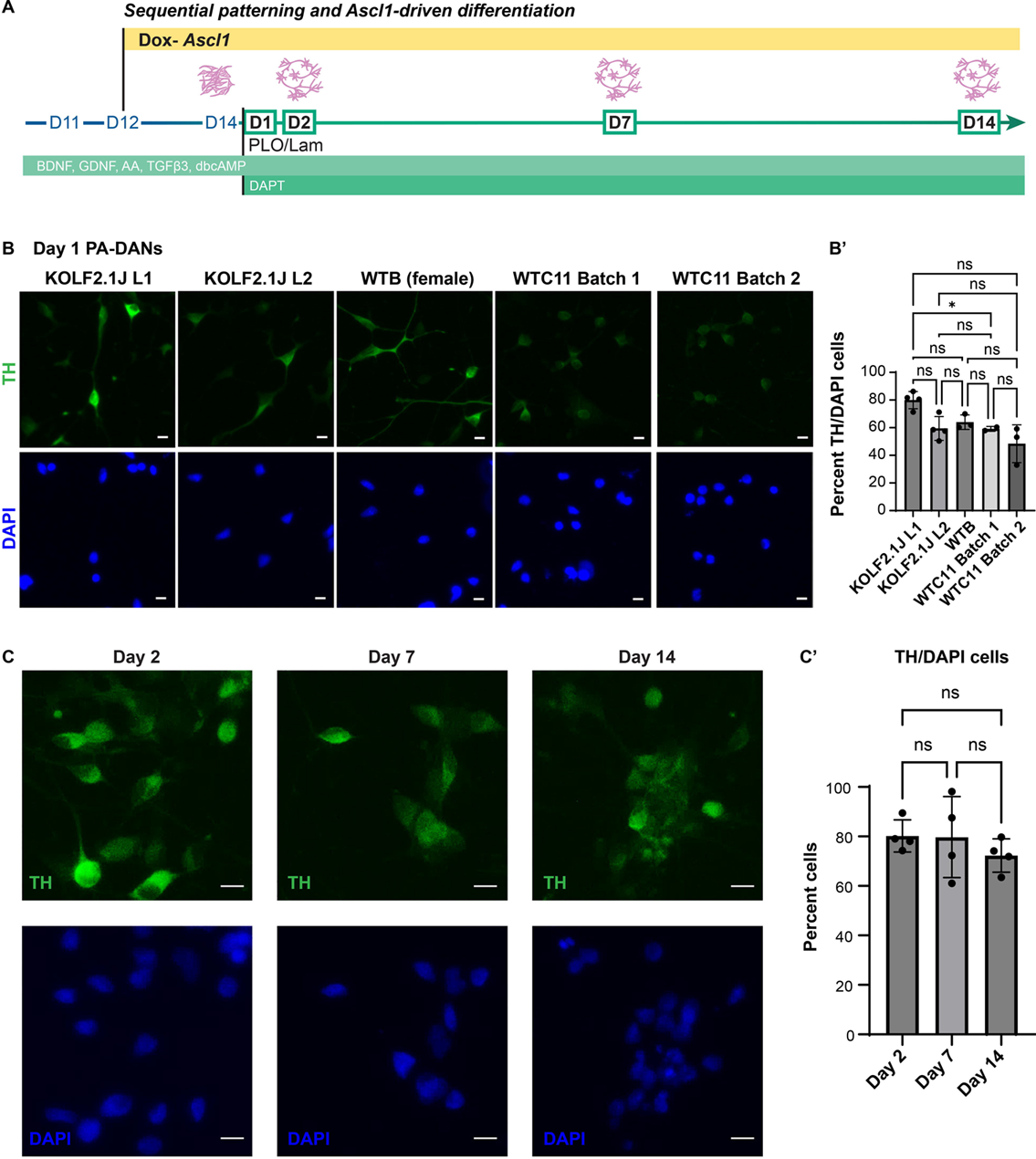
Ascl1 expression in patterned Ascl1-driven (PA-) progenitors leads to rapid differentiation of TH-positive dopaminergic neurons. A. Schematic showing the timeline of differentiation of PA-progenitors into PA-DANs. B. Representative images of PA-DANs differentiated for 24 hours and immunostained with TH (green) and co-stained with DAPI (blue). Scale bar 10µm. Lines tested: KOLF2.1J Line 1, KOLF2.1J Line 2, WTB, WTC11 batch 1 and WTC11 batch 2 (See methods). (B’) Graph showing high percentage of TH-positive PA-DANs after 24 hours of differentiating PA-progenitors generated via the sequential method. One-way ANOVA, multiple comparisons test, two tailed, *: >0.05, ns: not significant. Data are represented as mean ± SD. n=2-4, 2-3 fields from each replicate. Approx 2000 total cells counted per condition. C. Representative images of PA-DANs differentiated for 2, 8 and 14 days and immunostained with TH (green) and co-stained with DAPI (blue). Scale bar 10µm. (C’) Graph showing the percentage of PA-DANs expressing TH over days. One-way ANOVA, multiple comparisons test, two tailed, ns: not significant. Data are represented as mean ± SD. n=4, 3 fields from each replicate. Approx 2000 total cells counted per condition.

Next, we assessed the time course of DA neuron generation, by following the differentiation of DA neurons derived by sequential *Ascl1* expression in the presence of DOX, over several days (Figure 3A). We found that the percentage of cells expressing TH remained consistently at approximately 80% from Day 2 to Day 15 (Figure 3C, 3C’). We also immunostained these neurons after 24 hours of differentiation for additional markers such as SOX6, ALDH1A1 and Calb1 (Figure S1C). We found a high percentage of SOX6 and ALDH1A1 positive cells that represent the more ventral SNc (A9) population, and low percentages of Calb1 positive cells that correspond to the dorsal SNc (A9) and the ventral tegmental region (A10) (Figure S1C’). Interestingly, we also found that even within the first 18 hours of plating, DOX induction of Ascl1 with or without a Notch-inhibitor, DAPT, supported clear DA neuronal differentiation with two-fold higher percentage of TH-expressing cells (∼30% with DOX) than cells plated in the absence of DOX with (∼15%) or without DAPT (∼17%) which showed more progenitor-like morphology lacking processes (Figure S1D, S1D’). We termed the neurons and progenitors generated using the sequential *Ascl1* expression method as *Patterned Ascl1-driven DA Neurons (PA-DANs)* and *Patterned Ascl1-driven progenitors (PA-progenitors)*, respectively (Figure S1B). Taken together, we identified a crucial window during early iPSC-derived midbrain development wherein the expression of *Ascl1* could enhance DA neurogenesis.

### Single-cell RNA-sequencing of PA-DANS reveal midbrain specificity

To further confirm midbrain specificity and further elucidate the DA subtype enrichment, we characterized the PA-DANs during differentiation at days 1 and 15 using single-cell RNA-sequencing (Figure 4A). To assess the extent of terminal differentiation on Day 1, we identified two subgroups based on the expression pattern of *CCN1*, a cell cycle gene – roughly 20% of the cells expressing *CCN1* were classified as the progenitor subgroup, and ∼80% of the cells lacking *CCN1* expression were deemed terminally differentiated post cell cycle exit (Figure 4B, 4C). This group showed an enrichment of cells expressing *TH* mRNA, thus classified as the induced DA neuronal subgroup (Figure 4B, 4C). We observed expression of early midbrain markers such as *OTX2*, *FOXA2* and *LMX1a* across both the subgroups, confirming midbrain specificity, while other cell cycle genes such as *TOP2A* and *CDK1* were restricted to the progenitor subgroup and neural specific markers such as *TH* and *SNAP25* expressed more highly in the clusters corresponding to the induced DA subgroup (Figure 4C, 4D, Figure S3A, S3B). While expression levels of forebrain (*PAX6*) and hindbrain (*GBX2*) markers were either low or lacking, we found moderate levels of IRX3 known to be expressed in regions of both midbrain and hindbrain, according to the Human Protein Atlas (proteinatlas.org) (Sjöstedt et al., 2019), demonstrating that our protocol specified midbrain-restricted lineage (Figure S3A, S3B). We also observed a lack of other glial markers such as GFAP and OLIG2, indicating neuronal specificity. Importantly, we also observed high Ascl1 expression in the induced DA subgroup, reiterating the role of Ascl1 in cell cycle exit and DA differentiation (Figure S3A, S3B).

**Figure 4:**
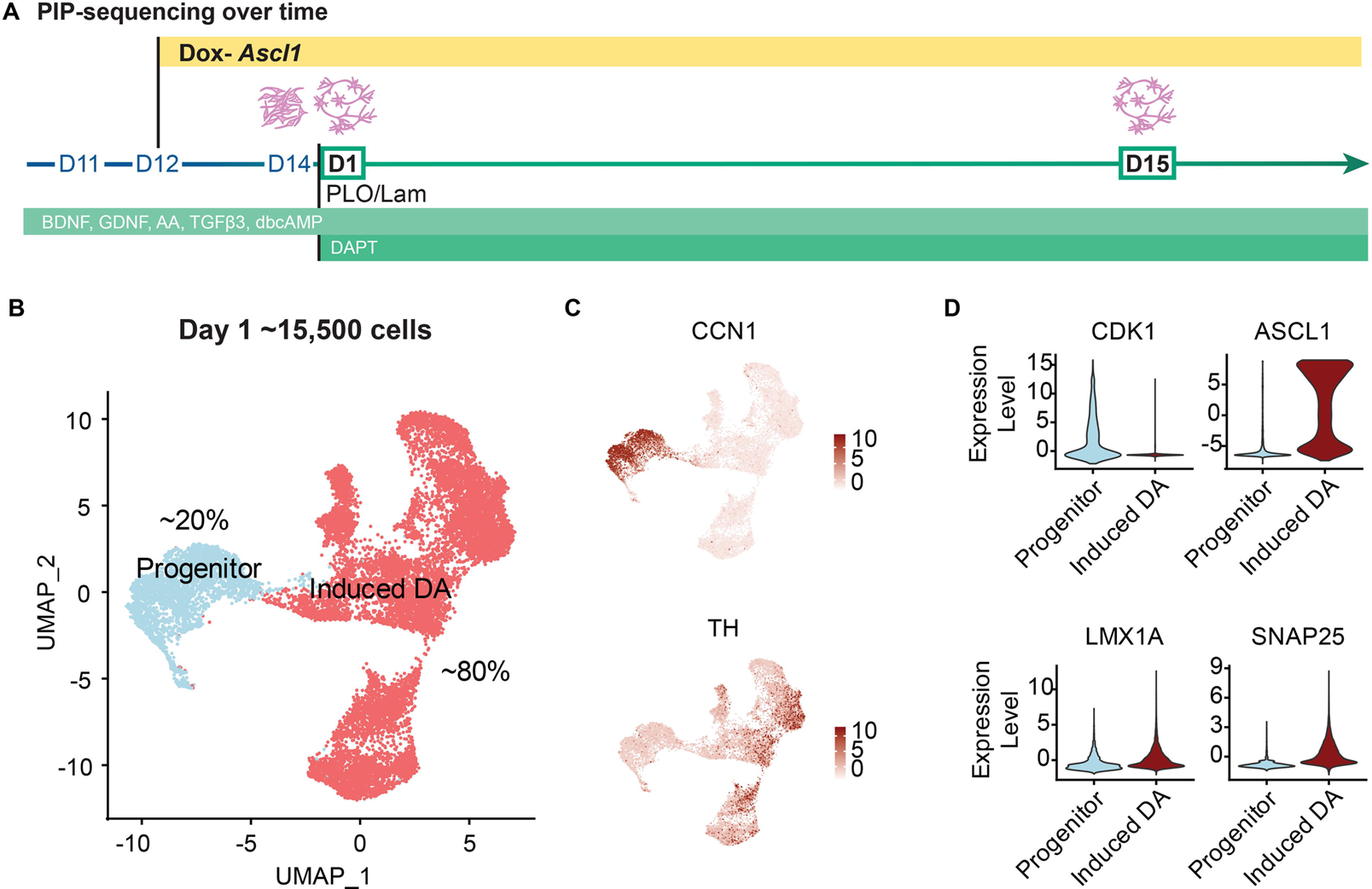
Transcriptomic analysis of Day 1 PA-DANs shows rapid terminal differentiation and midbrain specification with Ascl1 expression. See also Figures S2, S3. A. Schematic of PIP-Seq performed at Day 1 and Day 15 during differentiation. B. UMAP representation of Day 1 PA-DANs showing a distribution of progenitor-like and terminally differentiated induced DA-like subgroups. C. Feature plots showing enrichment of CCN1-positive or TH-positive populations D. Violin plots for key genes CDK1, Ascl1, LMX1a, SNAP25

Having established midbrain specificity and terminal differentiation at Day 1, we further characterized the extent of neuronal differentiation in PA-DANs at Day 15. Unsupervised clustering and visualization of Day 15 PA-DANs revealed a small progenitor population expressing *CDK1* and a terminally differentiated population lacking *CDK1* expression, which was enriched in *TH* mRNA (Figure 5A, 5B). To further explore the potential cell type heterogeneity in cells differentiated at Day 15, we employed a relatively high Principle Component (PC) number (JackStraw procedure, see methods) and identified 16 clusters (Figure S4A). Based on gene expression we classified these clusters into the progenitor group marked by expression of *CDK1*, and the differentiated group further divided into classes 1, 2 and 3 (Figure S4A, S4B). We first confirmed terminal neuronal differentiation across the different classes based on the expression of neuron-specific markers such as *SNCA* and *STMN2* – where clusters in Class 1 showed low expression, and clusters in Classes 2 and 3 showed high expression (Figure S4B). Conversely, we found negligible expression of astrocytic (*S100β*) and oligodendrocytic (*SOX10*) markers, indicating the absence of glial differentiation (Figure S4B). Next, we investigated the presence of broad neuronal cell types in our Day 15 PA-DANs population. Based on low expression of genes such as *GLS*, *GRIN1* and *SLC17A7*, we concluded that glutamatergic neurons were absent in our Day 15 PA-DANs population (Figure S4B). We also confirmed the absence of striatal GABAergic neurons based on the sparse expression of *PPP1R1B* (Figure S4B). On the other hand, we found a moderate to high expression of DA markers such as *TH*, *DDC* and *KCNJ6* as well as GABA receptors and receptor associated proteins, *GABARAP* and *ABAT*, across clusters, typical of A9-DA subtypes (Figure S4B) (Kamath et al., 2022). Finally, we compared the average expression levels of a list of 22 midbrain and DA-associated genes against the expression levels of unbiased set of control genes in our cell population, thereby deriving a midbrain module score. We found that the average expression of DA-specific genes was significantly higher than the background control expression levels in approximately 66% of the cells at Day 15, confirming the persistence of relevant markers during differentiation in PA-DAN monocultures even in the absence of a supporting glial layer (Figure S4D). Overall, these data confirmed midbrain specificity of our differentiated DA neurons.

**Figure 5:**
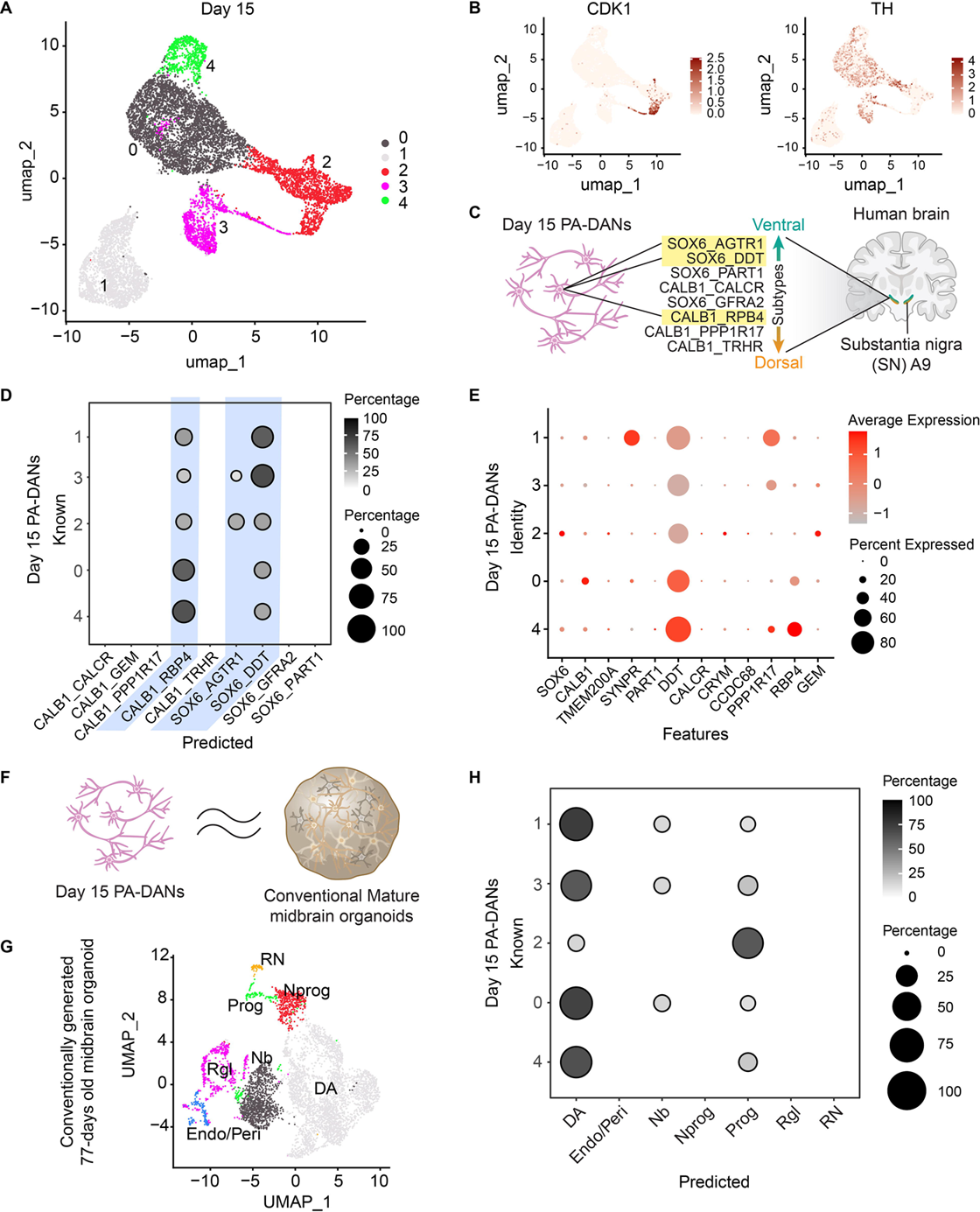
High correspondence of Day 15 PA-DANs with human A9-substantia nigra DA subtypes and DA neurons in conventionally generated midbrain organoids, based on machine learning algorithm. See also Figure S2, S4. A. UMAP representation of Day 15 PA-DANs showing 5 clusters B. Feature plot showing the enrichment of CDK1-positive and TH-positive populations. C. Schematic of DA subtypes in the human A9 region enriched in PA-DANs D. Dot blot (Confusion matrix) showing an enrichment of SOX6_DDT and CALB1_RBP4 DA subtypes. Small enrichment of the most vulnerable SOX6_AGTR1 DA subtype based on a trained machine learning algorithm E. Dot plot showing the expression of key genes associated with enriched subtypes F. Schematic depicting similarities of Day 15 PA-DANs to DA neurons in conventional midbrain organoids G. UMAP representation of 77 days old conventionally generated midbrain organoid (Kim et al, 2021) showing the distribution of different cell states including DA neurons, endocytes/pericytes (Endo/Peri), neuroblasts (Nb), Neural progenitors (Nprog), Progenitor cell types (Prog), Radial glia-like cells (Rgl), Red nucleus (RN) H. Dot blot (Confusion matrix) showing the high correspondence of Day 15 PA-DANs with DA neuronal cluster present in 77 days old conventional midbrain organoid determined via a trained machine learning algorithm.

### Single-cell transcriptomics show high correspondence with vulnerable DA subtypes of the ventral A9 region

To establish the DA neuronal subtypes enriched in the PA-DANs generated via our sequential method, we compared our dataset with existing adult human A9-SNc subtypes identified in a recent study (Kamath et al., 2022) (Figure 5C). To this end, we implemented an xgboost-based classifier as previously described (Chen and Guestrin, 2016; Hahn et al., 2023; Nimkar et al., 2025; Peng et al., 2019; Wang et al., 2024; Zhang et al., 2024) using the published adult human DA neuron dataset (training), to predict the label of each cell in our dataset (test). Briefly, the shared highly variable genes (HVGs) between the training and test datasets were identified as features, and the log-normalized expression matrices were then subset to these shared HVGs as training and test matrices. 60% of the training data was used to establish the model, followed by accuracy validation using the remaining 40% as a test. After validating a high accuracy, the model was then used to predict a label for each test cell based on the training category (See Methods). Using this framework, we found that the Day 15 PA-DANs correspond prominently to two A9-DA subtypes, SOX6_DDT and CALB1_RBP4 subtype (Figure 5D). These human DA subtypes correspond to the ventral and dorsoventral A9 region previously identified using the Macaque slide-sequencing of the SNc (Kamath et al., 2022). We found high DDT expression in our PA-DANs similar to that found in the SOX6_DDT and CALB1_RBP4 A9-DA subtypes (Figure 5E). In addition, we also see correspondence with a SOX6_AGTR1 subtype enriched in the more immature neuronal and progenitor-dominant clusters of PA-DANs (Figure 5D). This DA subtype corresponds to the most vulnerable subtype identified in human PD patients (Kamath et al., 2022). We confirmed high expression of key PD causative genes in our dataset (Figure S4C). Overall, we found that the subtypes enriched in our PA-DANs are in a continuum similar to the spatially overlapping DA subtypes in the human A9 region.

### Single-cell transcriptomics of PA-DANs show high resemblance to mature iPSC-derived DA neurons in midbrain organoids

To benchmark our approach against established methods for generating midbrain dopaminergic neurons, we further compared our Day 15 PA-DANs data to the published scRNA-seq dataset of Day 77 midbrain organoids generated using directed differentiation (Kim et al., 2024) (Figure 5F). To this end, we first obtained the accompanying expression matrix from GEO and classified the data by re-clustering and a compiled list of marker genes (Figure 5G). We then trained a classifier using the Day 77 midbrain organoid dataset based on the machine learning framework as mentioned above. This predicts a label for each cell in our Day 15 PA-DANs based on the Day 77 organoid data (See methods). We found that the cluster 2 in our Day 15 PA-DANs corresponded to both the progenitor and DA populations of the Day 77 midbrain organoids, while all the remaining clusters mapped prominently with the DA population, with a minor correspondence with neuroblast clusters (Figure 5H). Taken together, these results indicate that 2D monocultures of Day 15 PA-DANs generated via our sequential method are similar to DA neurons developed *in vitro* in older 3D midbrain organoid models.

### 3D assembled A9-like midbrain organoids show microglial integration and mature neuronal activity

We have previously shown that 3D organoids assembled using defined ratios of iNeurons, iAstrocytes and iMicroglia undergo greater cell type maturation in 3D as compared to 2D co-cultures (Li et al., 2025; Majo et al., 2023). Additionally, microglia integration into midbrain organoids is complicated given the lack of compatibility between DA differentiation factors and microglial viability as previously demonstrated and overcome in (Sabate-Soler et al., 2022). Here, we utilized our approach to generate 3D assembled A9-like midbrain organoids by combining Day 14 PA-progenitors along with iPSC-derived astrocyte precursor cells (iAPCs) in a 10:1 ratio representative of the human midbrain, in midbrain differentiation medium containing DOX (Figure 6A). We introduced iPSC-derived hematopoietic precursor cells (iHPCs) after one week of culturing the 3D assembled organoids to allow for microglial integration and differentiation (Figure 6A). To validate the maturation of all the cell types, we performed immunostaining for cell type-specific markers and found that 5 weeks old 3D A9-like assembled organoids show mature PA-DANs that express TH, MAP2 and VMAT2 (Figure 6B and 6C), mature GFAP-positive astrocytes and successfully integrated Iba1-positive microglia (Figure 6D). Microglia showed complex morphology typical of surveilling phenotype juxtaposed around neuronal cell bodies and axons (Figure 6D, Figure S5B). We also confirmed TH expression in differentiated PA-DANs in 4-week- old 3D assembled A9-midbrain organoids generated using PA-progenitors derived from a KOLF2.1J iPSC line containing a TdTomato reporter expressed under a TH promotor engineered using the CRISPR knock-in technology (Ahfeldt et al., 2020; Sarrafha et al., 2021). We observed positive immunostaining of anti-TH and anti-TdTomato antibodies in cells showing TdTomato fluorescence (Figure S5A). We further confirmed that these 3D A9-like assembled organoids retained midbrain specific expression of markers. Transcription factors such as FOXA2, Nurr1, LMX1A/B and SOX6 showed nuclear localization in 3D PA-DANs (Figure 7A, 7B, Figure S5C), similar to that seen in 2D cultures. We also observed low Calb1 intensity upon immunostaining in 3D as seen in 2D cultures of PA-DANs (Figure 7A). Further, we confirmed the expression of other A9-specific markers of PD-sensitive DA neurons, such as AGTR1 and ALDH1A1 in 5 weeks old A9-like assembled organoids (Figure 7B). In summary, we were able to generate a platform to study A9-like DA neuron and DA neuron-glia connections with distinct control over cell number and complexity.

**Figure 6:**
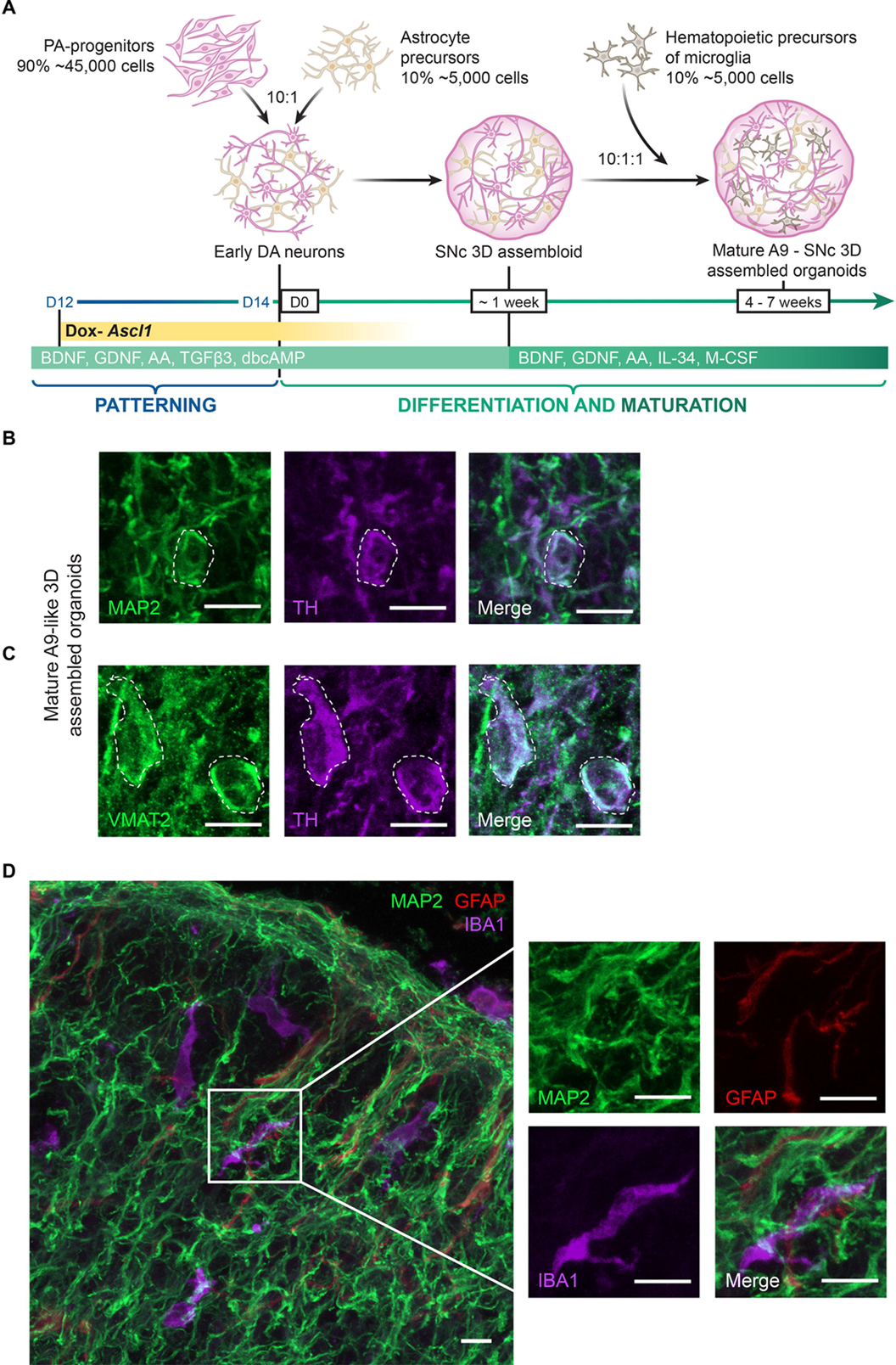
Generation of A9-like midbrain assembled organoids using PA-DANs. See also Figures S5, S6. A. Schematic showing the timeline for the generation of A9-like 3D organoids first assembled using PA-progenitors and iPSC-derived Astrocyte precursor cells (iAstrocytes) at a 10:1 ratio (50,000 total cells), followed by the addition of 5000 iPSC-derived hematopoietic precursor cells (iHPCs) after one week of co-differentiation and maturation in defined ratios reported in the human substantia nigra. B. Representative confocal images of 5 weeks old 3D organoids showing an overlap of MAP2 (green) and TH (purple) immunostaining of PA-DANs. Cell body is highlighted with broken white line boundary. Scale bar 10µm C. Representative confocal images of 5 weeks old 3D organoids showing an overlap of punctate VMAT2 (green) and TH (purple) immunostaining of PA-DANs. Cell bodies are highlighted with broken white line boundary. Scale bar 10µm D. Representative confocal images of 5 weeks old 3D organoids showing immunostaining against MAP2 (green), GFAP (red) and IBA1 (purple), visualizing PA-DANs, iAstrocytes and iPSC-derived microglia, respectively. Inset showing 2x magnified field of each cell type surrounding each other. Scale bar 10µm

**Figure 7:**
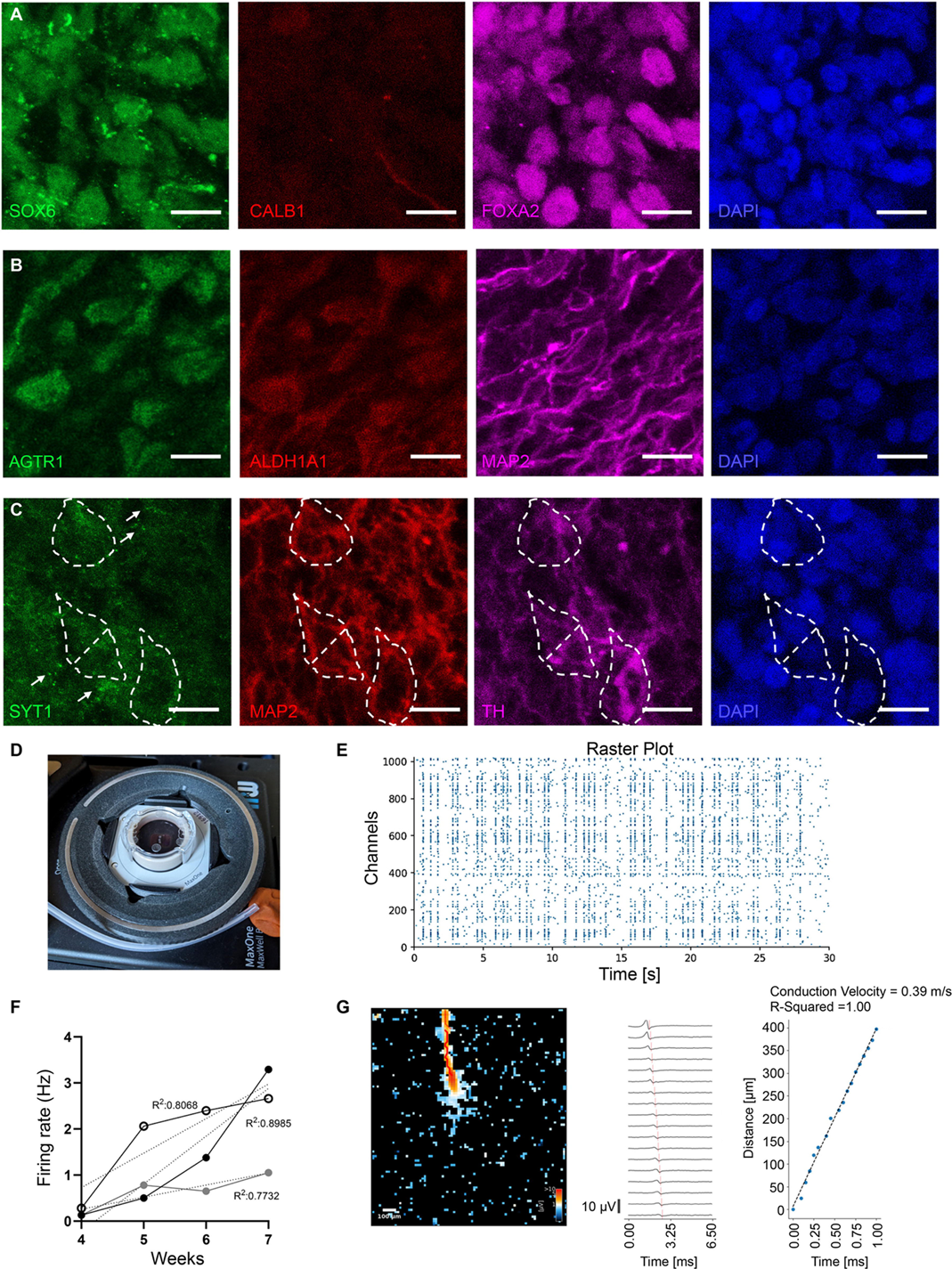
PA-DANs in A9-like 3D assembled organoids express A9-specific markers of selective vulnerability and show neuronal activity. A. Representative images of 5-weeks old A9-like 3D assembled organoids showing nuclear localization of ventral A9 marker SOX6 (green), midbrain marker, FOXA2 (magenta), cytoplasmic localization of dorsal A9 and A10 marker CALB1 (red), and nuclear marker DAPI (blue). Scale bar 10µm B. Representative images of 5-weeks old A9-like 3D assembled organoids showing perinuclear localization of A9 markers, AGTR1 (green) and ALDH1A1 (red), dendritic localization of neuronal marker MAP2 (magenta) and nuclear marker DAPI (blue). Scale bar 10µm C. Representative images of 5-weeks old A9-like 3D assembled organoids showing punctate localization of membrane-associated protein, SYT1 (green), dendritic localization of neuronal marker MAP2 (red), TH (magenta) and nuclear marker DAPI (blue). Cell bodies are highlighted with broken white line boundaries. Scale bar 10µm D. Image showing MEA setup with three 4-weeks old A9-like 3D assembled organoids plated on a MaxOne chip (MaxWell Biosystems AG) E. Increasing firing rate of neurons in A9 over time showing 1-3 Hz firing typical of midbrain DA neurons from 3 independent experiments with non-significant difference between their slopes (P=0.1068). Measurements with MEAs (MaxWell Biosystems AG) F. Example of a Raster plot showing periodic activity of PA-DANs in 7-weeks old A9-like 3D assembled organoids. Measurements with MEAs (MaxWell Biosystems AG) G. Example of a single axonal firing and propagation with a conduction velocity of 0.39 m/s of PA-DANs in 7-weeks old A9-like 3D assembled organoids. Measurements with MEAs (MaxWell Biosystems AG)

### Multielectrode Array (MEA) recordings show spontaneous firing and network activity in patterned *Ascl1*-driven dopaminergic neurons in 3D assembled A9-like organoids

To assess the functionality of PA-DANs in the 3D environment of assembled A9-like organoids, we performed MEA recordings to measure the firing pattern of population spikes and axonal conductance from individual cells (Figure 7D). We found that the spontaneous firing rate steadily increased from 0.5Hz to 3Hz in 4 to 7 weeks of culturing 3D A9-like assembled organoids on MEA chips. The firing pattern shows a typical tonic firing, which is reported previously in DA neurons in the SNc and in *in vitro* DA models (Figure 7E) (Grace et al., 2007; Ronchi et al., 2021; Shi, 2009). The neurons showed synchronous firing in network at periodic intervals, with an interspike interval centered around ∼100ms (Figure 7F) (Nolbrant et al., 2024). This indicates the neurons in the 3D A9 assembled organoids form connections among each other and fire together. Further, we measured the axon conductance velocity of individual DA neurons in 3D A9-like assembled organoids and found it to be ranging between 0.4 – 0.7 m/s, which was similar to the range that has been previously reported (Figure 7G) (Ronchi et al., 2021; Shi, 2009). Concurrently, we found clear labelling of VMAT2 (Figure 6C) and Syt1 (Figure 7C) by immunostaining, necessary for dopamine release (Alwindi and Bizanti, 2023; Delignat-Lavaud et al., 2023; Erickson et al., 1996; Lebowitz et al., 2022). Taken together, these analyses confirmed that PA-DANs mature and exhibit DA activity in 3D A9-like assembled organoids.

## Discussion

The substantia nigra pars compacta (A9) consists of heterogenic subtypes of DA neurons distributed dorsoventrally in the midbrain, wherein the ventral SOX6-positive DA subtypes are more vulnerable (Kamath et al., 2022; Luppi et al., 2021; Poulin et al., 2014). To date, published protocols fail to generate subregional-specific DA subtypes corresponding to the SNc. In this study, we enlisted the use of transcription factor expression into the directed differentiation approach. Here we report, a method to rapidly derive PD-sensitive subtypes of DA neurons from human iPSCs; by sequentially using developmental cues to pattern floor plate mesencephalic progenitors *to ensure midbrain specificity,* and the expression of a neurogenic transcription factor, *Ascl1*, expressed in the midbrain lineage and known *to enhance DA neuronal differentiation*. We found that the timing of Ascl1 expression largely influenced the quality and homogeneity of the resulting neuronal cultures. We chose to induce Ascl1 expression in early floor plate mesencephalic progenitors based on the known temporal role of Ascl1 during early midbrain development in human and animal studies. Our method extends the concepts explored in a very recent study investigating the use of neurogenic TF expression in post-patterned iPSCs to generate a variety of broadly different, regionally and functionally defined neuronal subtypes (Lin et al., 2025). In our study, we have refined this strategy to examine DA subregional heterogeneity and developed ventral A9-like DA neurons crucial in studying Parkinson’s disease in vitro.

The timing of Ascl1 expression over the course of midbrain development is crucial. Ascl1 expression starts in cycling progenitor and continues through transition states into terminally differentiated neurons. Ascl1 works by regulating cell division and inhibiting cell cycle genes, thereby helping progenitors exit cell cycle (Altbürger et al., 2023). These studies support the result that induction of Ascl1 on Day 12, rather than on Day 1, during the early floor plate mesencephalic progenitor development from iPSCs, enhances DA differentiation. Ascl1 expression is also regulated via Notch inhibition during development (Park et al., 2017), inducing neurogenesis, which we observed in our PA-DANs, where 18 hours of differentiation in the presence of Dox and absence of DAPT was sufficient for robust neurogenesis. Ascl1 is a non-canonical pioneer factor working with other chromatin machinery (mSWI/SNF) making it more versatile depending on the cell state (Păun et al., 2023). Loss of Ascl1 expression impedes neurogenesis, accumulating cycling progenitors, even in the presence of DAPT (Lundie-Brown et al., 2025). We made a similar observation where in the absence of inducible Ascl1 expression, the early floor plate mesencephalic progenitors failed to differentiate after 18 hours of exposure to neurogenic factors and DAPT. In our transcriptomic analysis, we found that cell cycle genes are downregulated in Ascl1 -high expressing cells at Day 1. This implied that inducing Ascl1 expression ensures a more congruent transition of progenitors into terminally differentiated neurons.

We chose to utilize Ascl1 as opposed to NGN2 for several reasons. Ascl1 strongly initiates and regulates GABAergic differentiation, whereas NGN2 strongly supports glutamatergic differentiation (Parras et al., 2002). Through meta-analysis, we found that in the human fetal midbrain transcriptome, Ascl1 expression is prominent in the midbrain progenitors, but also continues in DA and GABAergic neuronal clusters. NGN2 expression remains lower overall in the human fetal midbrain dataset but it is also expressed in DA neurons (Data not shown) (Braun et al., 2023; Nolbrant et al., 2024). While both Ascl1 and NGN2 play a role in dopaminergic differentiation, Ascl1 expression can partially rescue midbrain development in mice lacking NGN2 (Kele et al., 2006; Kim et al., 2007). Ascl1 has been shown to upregulate TH in eSCs and ventral mesencephalic primary neuronal precursor cells as opposed to NGN2 (Kim et al., 2007; Ng et al., 2021). Previously, Ascl1 expression in combination with other midbrain-specific transcription factors such as Nurr1 and LMX1 has been used in iPSC-derived dopaminergic specification (Ng et al., 2021; Powell et al., 2021; Theka et al., 2013). Moreover, exogenously expressed Ascl1 has been shown to transdifferentiate midbrain astrocytes into neurons in vivo, further underscoring the potency of Ascl1 in DA neurogenesis (Liu et al., 2015; Yong et al., 2024). Overall, our data is consistent with the hypothesis that Ascl1 is sufficient to drive ventral A9 regional specificity to early floor plate mesencephalic progenitors in vitro.

Our findings extend recent efforts to refine directed differentiation through temporally coordinated transcription factor induction.to not only enhance neurogenesis but also promote subtype identity. Recent protocols in developing regionally-specific neurons have fine-tuned directed differentiation with concurrent NGN2 expression, such as generating more regionalized and mature NGN2-derived cortical and motor neurons (Limone et al., 2023; Nehme et al., 2018; Shan et al., 2024). In another strategy, NGN2-induced neural progenitors, rather than iPSCs, have been differentiated in DA neurons using commercially available midbrain differentiation kit (McDonald et al., 2024; Sheta et al., 2022). Most recently, a study that screened for over 400 combinations of small molecule morphogens and TFs has similarly concluded as our study that sequential early patterning followed by TF expression yields more consistent differentiation and continued maturation (Lin et al., 2025). Through our work, we postulate an additional application of this premise - that optimizing early patterning to produce regional progenitors and identifying transcription factors that can further drive subregional specification are the keys to generating dopaminergic neuronal subtypes.

We have additionally pioneered a rapid and robust method to generate active 3D assembled organoids using human iPSC-derived cortical-like neurons and astrocytes having precise cellular ratios thereby removing previous limitations such as lack of control over cell maturity and number (Li et al., 2025; Majo et al., 2023). We have modified this technique to develop A9-specific 3D midbrain assembled organoids containing only PA-DANs, midbrain iPSC-derived astrocytes, and iPSC-derived microglia in precise numbers that reflect the ratios present in the human midbrain, unlike current midbrain organoid methods. Transcriptomic analysis of current models of human midbrain organoids and iPSC-derived DA neuronal populations contain cell types such as radial glial cells, progenitors and DA neurons, and occasionally glial cells, in different stages of development attributing to heterogenous differentiation (Fiorenzano et al., 2021). These often do not represent the human midbrain cell constitution. Importantly, successful human iPSC-derived microglial integration into the 3D midbrain organoids has only been reported in one other recently published *in vitro* model for human midbrain (Sabate-Soler et al., 2022). Microglia integration into midbrain organoids is challenging and it is difficult to replicate the complex ramified in vivo microglial maturation and morphology in vitro (Augusto-Oliveira et al., 2025). Our unique approach of assembling mature 3D midbrain organoids containing A9-like DA neurons with glial cells serve as an excellent model to study intricate midbrain-specific cell-autonomous and non-cell-autonomous mechanisms in aging and disease progression.

### Limitations of the study

We describe here a rapid method to generate a high yield of A9-like DA neurons from iPSC by using a sequential approach combining small molecules and transcription factor induction. We have characterized our PA-DANs in 2D monocultures using immunostaining and single-cell transcriptomics in the absence of a supporting astrocytic layer. While we observe neuronal morphology and expression of nigral DA neuronal genes, the cells show low expression of neuronal maturation such as VMAT2 and DAT over longer periods of time in 2D monocultures. Most published directed differentiation methods do not include external supporting astrocytes for DA maturation as these long-term cultures are supported by slowly differentiating progenitors and small percentage of intrinsic glial populations. However, some versions of TF induction methods for stem cell derived DA generation use astrocytes that bolster DA maturation, while many other versions do not. Given the combinatorial effect of directed and TF induced differentiation protocols in our method, addition of glia in 2D PA-DAN cultures is expected to assist further maturation, similar to what we observe in 3D assembled organoids in our system. Secondly, while we see enrichment of two or three ventral and dorsoventral DA subtypes in Day 15 PA-DANs using our sequential method, further perturbation of the protocol will likely achieve more precise subtype identity, specifically the vulnerable subpopulations in PD. Thirdly, our A9-like 3D assembled midbrain organoids generated using these PA-DANs show markers of PD sensitive DA neurons, such as SOX6, AGTR1 and ALDH1A1. However, we lack control over the spatial organization of different cell types in the assembled organoid model which may also help improve the DA subtype specificity in 3D. Further refinement of our protocol would be useful in improving cell type specificity and reducing GABAergic population as well. Overall, our method generates PA-DANs as modular components to assemble organoids with glial cell types and create 3D environments that support more mature morphologies and functionality for both DA neurons and glia.

## STAR Methods

### Key resource table

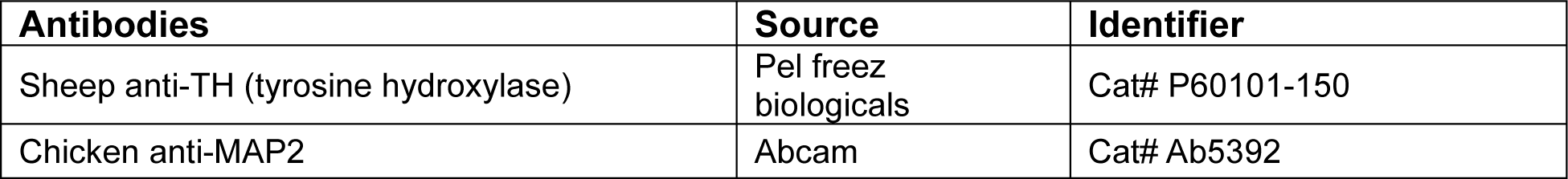
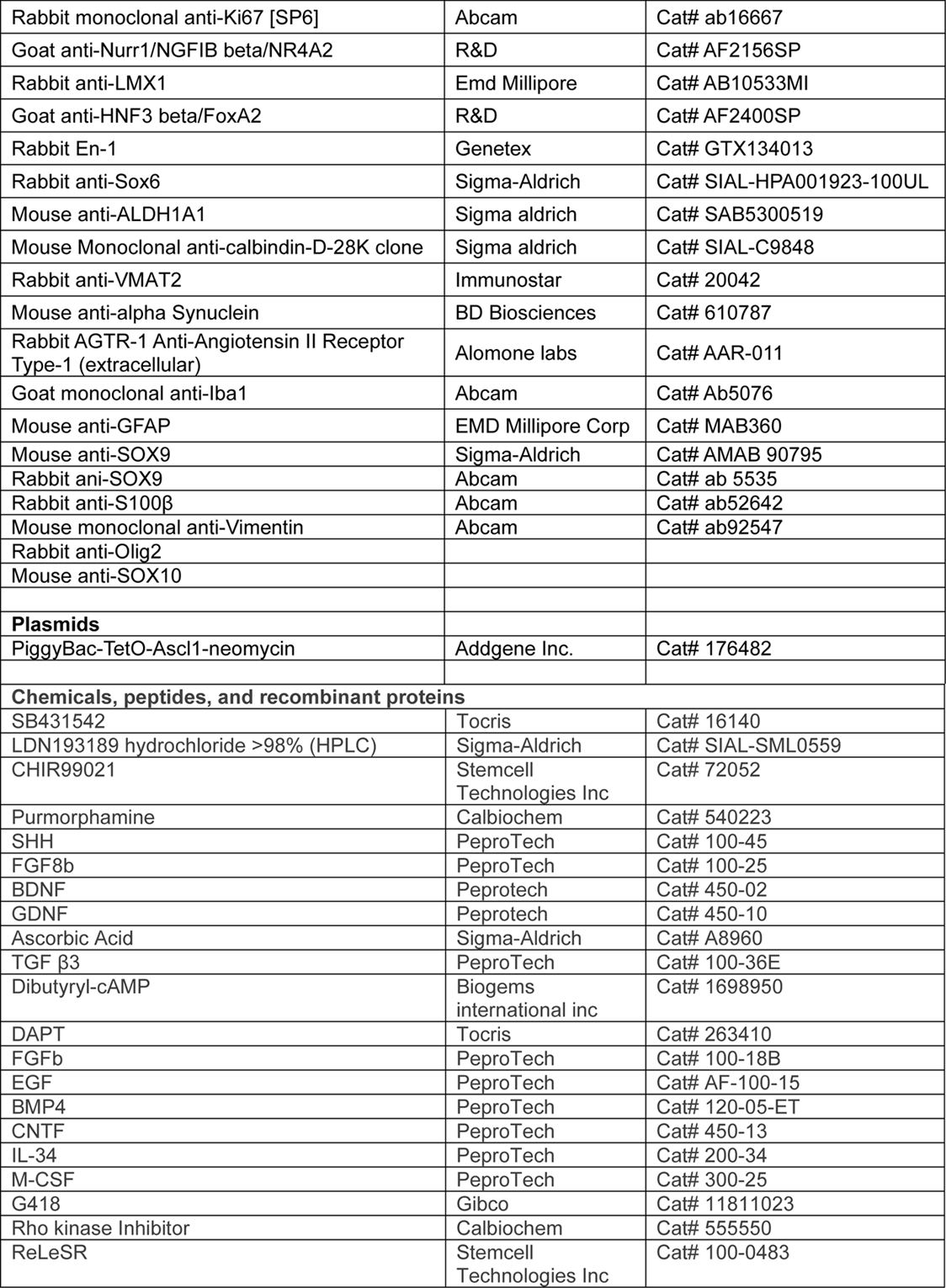
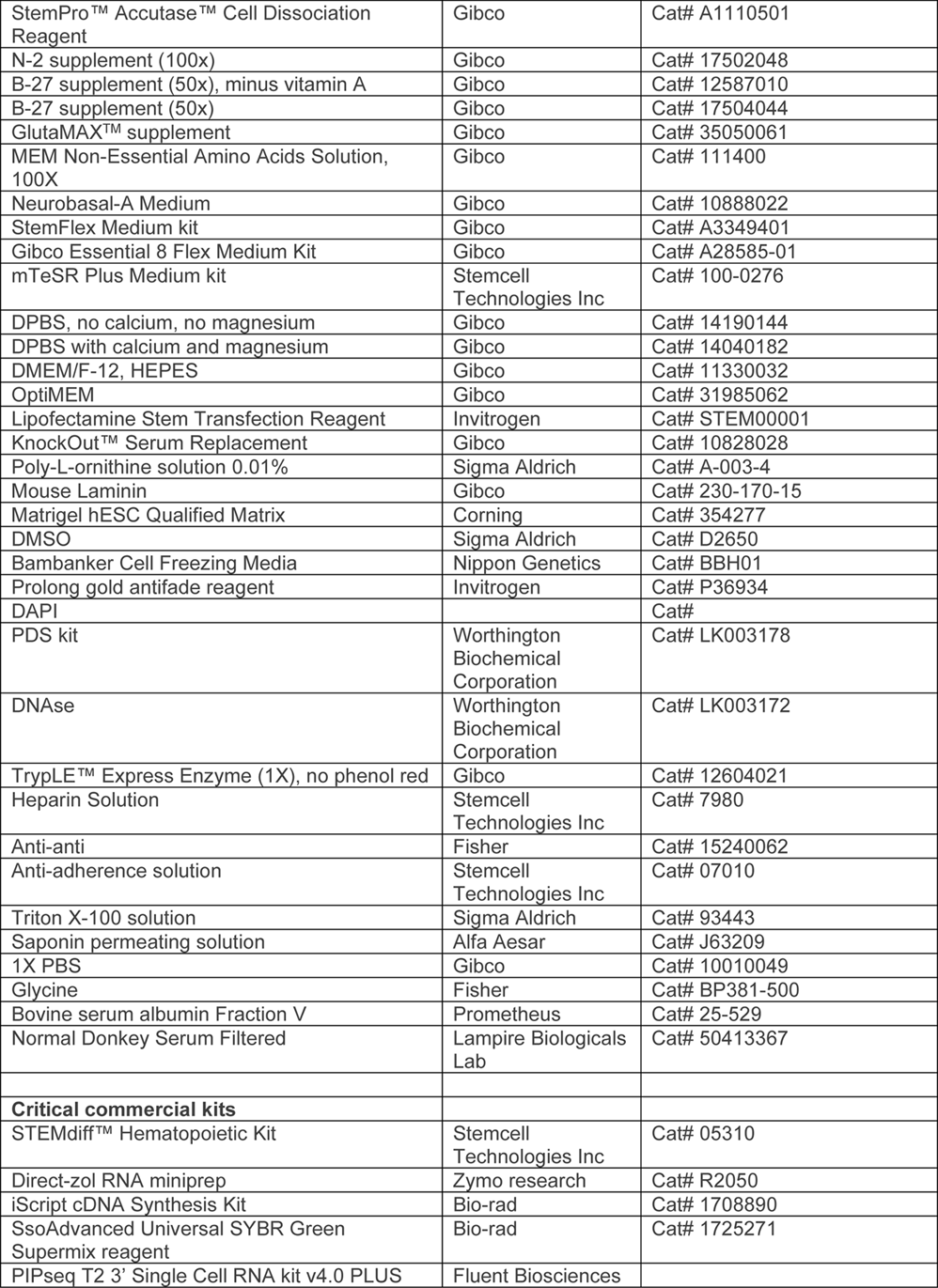
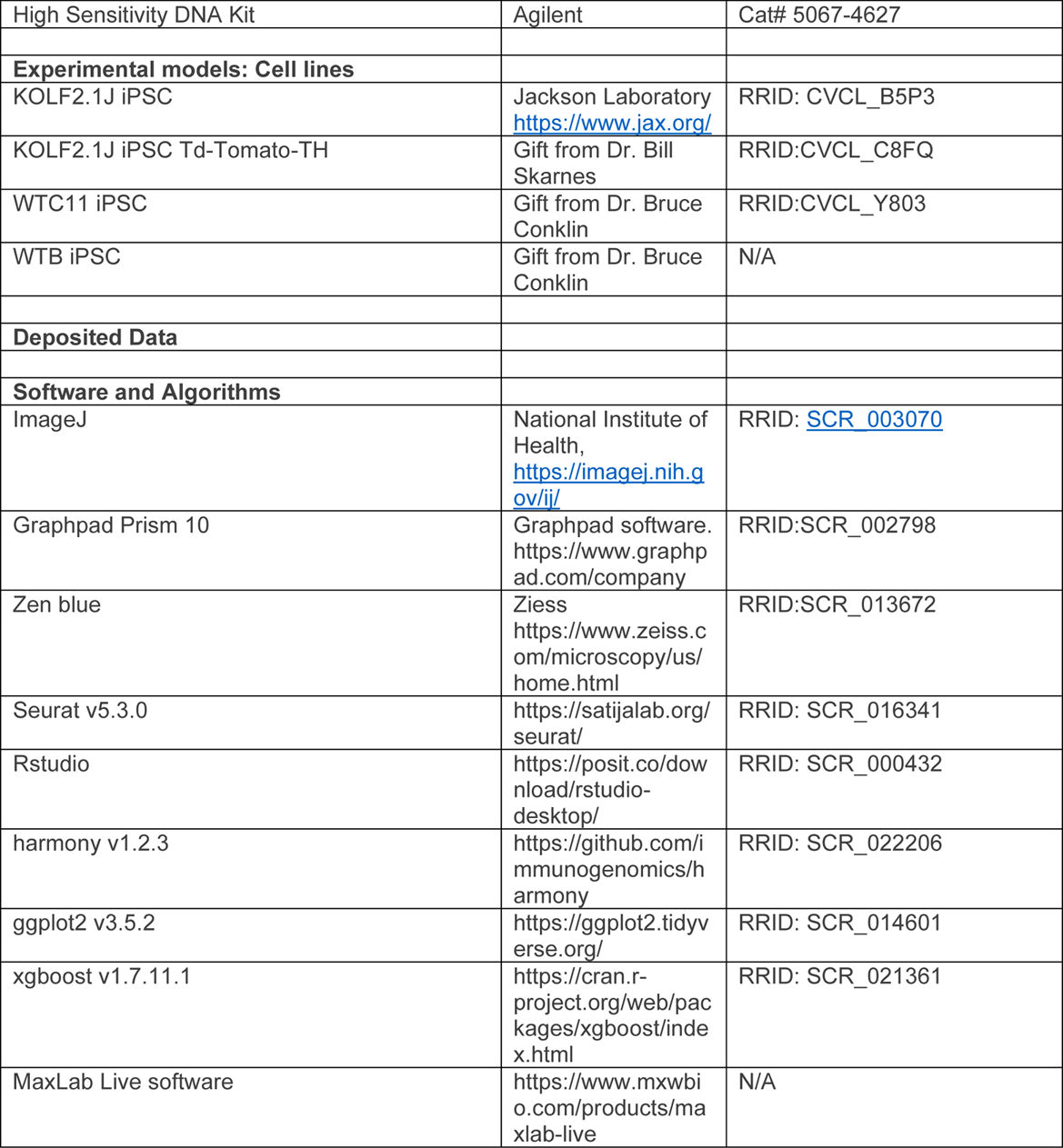

### Generation of Ascl1-piggybac iPSC line

The human KOLF2.1J iPSCs (male; Jackson Labs; named KOLF2.1J Line 0 and used throughout the study for all experiments unless otherwise mentioned) (Pantazis et al., 2022) and KOLF2.1J iPSCs containing TH promoter-driven Tdtomato (male; Gift from Dr. Bill Skarnes) (Ahfeldt et al., 2020; Sarrafha et al., 2021), were cultured in StemFlex medium (Gibco; A3349401). WTB (female; Gift from Dr. Bruce Conklin) and WTC11 iPSCs (male; Gift from Dr. Bruce Conklin, Gladstone Institute), were cultured in mTeSr Plus (Stemcell technologies; 100-0276) and Essential 8 Flex medium (Gibco; A2858501), respectively. All cell lines were characterized for pluripotency markers and by standard karyotyping. Each line was transfected by a piggybac vector encoding the mouse ***Ascl1*** gene (PiggyBac-TetO-Ascl1-neomycin, Gift from Marius Wernig (Addgene plasmid # 176482; http://n2t.net/addgene:176482; RRID:Addgene_176482)), which is highly conserved in humans, and a vector encoding EF1alpha transposase at a 3:1 ratio (Ng et al., 2021) using the Lipofectamine Stem transfection reagent (Gibco; STEM00001) as per the manual’s instructions. The cells were selected and maintained in growth medium containing the antibiotic, G418, at the concentration of 50µg/ml for successful selection of iPSCs integrated with piggybac-Ascl1. We generated one piggy-bac transfected line from KOLF2.1J Line 0 (L0 – used for all experiments unless specified otherwise)), two piggy-bac transfected lines from KOLF2.1J TH promoter-driven Tdtomato (KOLF2.1J Line 1 in Figure 2, Figure 3B; and KOLF2.1J Line 2 in Figure 3B, Figure S5), one line from WTB (Figure 3B) and one line from WTC11 (Batch 1 PA-DANs in Figure 2; Batch 2 PA-DANs in Figure 2 and Figure 3B). All iPSC cell lines with rounded colonies at 70-80% confluency were passaged using ReLeSR (StemCell technologies; 100-0483). Frozen stocks were made in Knock-out serum replacement (KOSR) medium with 10% DMSO after 3 passages. Cells were not used and maintained beyond 15 passages to ensure stemcellness and piggybac-Ascl1 transposon integration. To generate Ascl1-driven neurons, iPSCs were plated at a density of 100k cells/ sq. cm and induced with doxycycline (Dox 2µg/ml) in NB(N2B27) - Neurobasal-A medium (Gibco; 10-888-022) containing 0.5x B27 supplement without Vitamin A (Gibco 12587010), 1x N2 supplement (Gibco; 1702048), 1x GlutaMAX (Gibco; 35050061), 1x MEM Non-Essential Amino Acids (NEAA) (Gibco; 11140050) and cell death inhibitor [10µM Y27632 dihydrochloride ROCK inhibitor (Tocris; 125410)]. Medium was replaced next day with NB(N2B27) containing Dox. Cells were harvested on Day 3 and frozen in Bambanker (Nippon Genetics; BB05).

### Generation of iPSC-derived floor cell mesencephalic progenitor cells

For midbrain specificity, we modified the previously described protocol (Calatayud and Muñoz-Pedrazo, 2022) used to generate floor plate mesencephalic progenitors. To pattern the iPSCs, on Day 0, the Ascl1-PiggyBac containing iPSCs were dissociated using Accutase (Innovative cell technologies; AT104) for 10 minutes at 37 deg Celsius and plated at a density of 400k cells/sq. cm/well in a 6 well plate in 3ml of NB(N2B27) - Neurobasal-A medium (Gibco; 10-888-022) containing 0.5x B27 supplement without Vitamin A (Gibco 12587010), 1x N2 supplement (Gibco; 1702048), 1x GlutaMAX (Gibco; 35050061), 1x MEM Non-Essential Amino Acids (NEAA) (Gibco; 11140050). **Patterning factors were added to this medium to modulate key midbrain-specific pathways: SMAD inhibitors** [10µM SB431542 (Tocris; 1614 -50mg) and 500nM LDN193189 hydrochloride (SIGMA; SML0559-5MG)]; **SHH pathway activators** [200ng/ml Human SHH protein (Peprotech; 100-45), 0.7µM Purmorphamine (SIGMA; 540223-5MG)]; **Wnt signaling activator** [0.7µM CHIR99021 (StemCell Technologies; 72054); and cell death inhibitor [10µM Y27632 dihydrochloride ROCK inhibitor (Tocris; 125410)]. Throughout the protocol, cells were rinsed with DPBS containing calcium and magnesium (Gibco; 14040141) before being fed with fresh medium. On Day 1, cells fed with the Day 0 medium without the ROCK inhibitor. For the **concurrent method**, doxycycline (Dox 2µg/ml) was also added to the medium on Day 1 (Figure 1A, panel i). On Day 3 and Day 5, cells were fed with the above described medium with the concentration of CHIR99021 increased to 3µM. On Day 7 and 8, SMAD inhibition and SHH activation were discontinued, the cells were fed with 4ml of NB(N2B27) with only 3µM CHIR99021 for continued Wnt activation. On Day 9, cells were fed with the above NB(N2B27) medium containing 3µM CHIR99021 and 100ng/ml Fibroblast Growth Factor 8b (FGF8b) for activate **Wnt and FGF8 signaling**, respectively. On Day 10, the cells were fed with NB(B27) - Neurobasal-A medium containing 0.5x B27 without vitamin A, 1x GlutaMAX – with **midbrain-specific neurotrophic and pro-survival factors** [20ng/ml Brain-Derived Neurotrophic Factor (BDNF), 20ng/ml Glial cell line-Derived Neurotrophic Factor (GDNF), 1ng/ml Transforming Growth Factor β3 (TGFβ3), 200µM dibutyryl-cAMP, 200µM Ascorbic acid, 3µM CHIR99021, 100ng/ml FGF8b] and cell death inhibitor [10µM Y27632 dihydrochloride ROCK inhibitor]. Next day, the cells were dissociated with Accutase for 10 minutes at 37 deg Celsius to produce a single cell suspension. The cells were plated at a density of 800k cells/sq. cm/well in a 6 well plate in the same medium composition as Day 10, except the Wnt activator, CHIR99021. For **sequential method**, on Day 12, the cells were rinsed and fed with Day 11 medium composition (without cell death inhibitor) containing 2µg/ml of **doxycycline to induce the expression of Ascl1** (Figure 1A, panel ii). Dox addition was omitted for patterning without Ascl1 expression (Figure S1A, panel ii). After 48 hours, the progenitor cells were dissociated using the Papain dissociation system (Worthington Biochemical Corporation) as described previously (Jerber et al., 2020). Briefly, cells were treated with a cocktail of accutase:1x DPBS (1:1) containing papain (Worthington Biochemical Corporation; LK003176) at 37°C for 12 minutes and quenched with NB(B27) medium containing ROCK inhibitor and DNAse D2 (Worthington Biochemical Corporation; LK003170). Cells were spun down at 300g for 5 minutes and frozen in NB(B27) + 10% DMSO in a cell freezing container in -80°C freezer and then moved to liquid nitrogen for long-term storage.

### Generation of iPSC-derived patterned Ascl1-driven dopaminergic neurons

For differentiation using the sequential method, 14 days old progenitors induced for Ascl1 expression were plated at a density of 100k cells/sq. cm in glass chamber slides, coated with 0.01% poly-L-ornithine (Sigma, A-004-C) and 1µg/ml mouse laminin (Gibco, 230-170-15), in NB(B27) medium containing **neurotrophic and maturation factors** [20ng/ml BDNF, 20ng/ml GDNF, 200µM Ascorbic acid, 1ng/ml TGFβ3, 200µM dibutyryl-cAMP]; **Notch pathway inhibitor** [10µM DAPT], Ascl1 inducer [2µg/ml doxycycline]; and cell death inhibitor [10µM Y27632 dihydrochloride ROCK inhibitor] (Figure 1A, panel ii). Next day, the cells were fed with the same medium as above without the cell death inhibitor. The developing neurons were fed every 3-4 days. Progenitors generated by expressing Ascl1 alone (Figure S1A, panel i), by patterning iPSCs for midbrain lineage (Figure S1A, panel ii) and by using the concurrent method (Figure 1A, panel ii), were similarly plated and cultured to test for cell morphology.

### Generation of iPSC-derived Astrocytes and microglia

The iPSC-derived astrocyte precursor cells (iAPCs) were generated based on the method described previously (Krencik and Zhang, 2011; Krencik et al., 2017; Majo et al., 2023). Briefly, KOLF2.1J Line 0 iPSCs were patterned for midbrain specificity (as described above) for a duration of 16 days to create rosettes that were selected on Day 11 and Day 16 using the STEMdiff Neural Rosette Selection reagent (STEMCELL technologies; 05832). The iAPCs were enriched and matured in medium containing EGF and FGF cytokines for 3-10 months. In 2D cultures, iAPCs were differentiated into mature astrocytes over 7 days with the addition of BMP4 (10ng/ml; Peprotech) and CNTF (10ng/ml; Peprotech) every two days. The KOLF2.1J iAstrocytes were characterized by immunostaining against astrocytic markers such as GFAP, SOX9 and S100beta (Figure S7). Previously published WTC11 iAPCs were use in Figure S5 (Krencik and Zhang, 2011; Krencik et al., 2017; Majo et al., 2023). The iPSC-derived hematopoietic precursor cells (iHPCs) were generated from iPSCs using the STEMdiff hematopoietic differentiation kit (STEMCELL technologies #05310) using the step-by-step protocol described previously (McQuade and Blurton-Jones, 2021). Briefly, iPSCs were harvested in 40-60 clusters of ∼100 cells each and plated in iPSC-feeding medium. Media containing components A or B were applied as previously indicated over the course of 10-12 days. Cells were collected on Days 10 and 12 and used directly or frozen stocks were made in Bambanker (Nippon Genetics; BBH01) until use.

### Generation of mature 3D midbrain assembled organoids

96-well V-shaped bottom plates (Costar; 3894) were pre-treated with anti-adherent solution (Stem Cell Technologies; 07010) right before cell seeding. Day 14 patterned Ascl-driven progenitors and iPSC-derived astrocyte precursor cells (APCs) were seeded (10:1 iDA:iA ratio; 50,000 cells total per well) in 100µl of NB(B27) medium containing neurotrophic factors [20ng/ml BDNF, 20ng/ml GDNF, 200µM Ascorbic acid, 1ng/ml TGFβ3, 200µM dibutyryl-cAMP]; Ascl1 inducer [2µg/ml doxycycline]; and cell death inhibitor [10µM Y27632 dihydrochloride ROCK inhibitor]. After 24 hours, an additional 100µl of NB(B27) medium containing the neurotrophic factors same as the day before, was added and the organoids were allowed to continue to aggregate. To generate triple-cell type 3D organoids, five - seven days after aggregation, approximately 5000 iPSC-derived hematopoietic precursor cells (HPCs) were added to integrate into the immature assembled organoids at a ratio of 10:1:1 iDA:iA:iM in NB(B27) containing 20ng/ml BDNF, 20ng/ml GDNF, 200µM Ascorbic acid. Two cytokines, IL-34 (100ng/ml) and mCSF (25ng/ml) were added to the medium to aid microglial differentiation. A 50% media change is performed every 5-7 days for continued maturation of 3D assembled organoids (Figure 6A).

### Immunostaining

2D cells were grown and differentiated in poly-L-ornithine and laminin-coated 4-chambered slides for the times reported. Immunostaining for 2D cell types was performed by first fixing them with 4% paraformaldehyde for 15-20 minutes, followed by 3 quick rinses with 1x PBS. Cells were blocked in 1x PBS buffer containing 5% normal donkey serum (NDS) and 0.5% saponin for one hour at room temperature. Cells were incubated overnight at 4 °C in primary antibodies diluted in blocking buffer. After 3 quick 1x PBS rinses, cells were incubated in secondary antibodies and DAPI (to stain the nucleus) diluted in 0.1% bovine serum albumin (BSA) solution made in 1x PBS for two hours at room temperature. For immunostaining 3D organoids, organoids were fixed in 4% paraformaldehyde (containing 0.1%Triton X-100) for 40-60 minutes, followed by three 1X PBS washes. Fixed organoids were embedded in OCT molds and cut into 20µm thick slices using Cryostat, collected on slides. Slides were stored at -80 °C until use. Slices were circled with hydrophobic pen on slides and blocked with 1x PBS buffer containing 5% NDS, 0.5% triton X- 100, 1% BSA and 0.1M glycine for 1 hour. As performed for 2D cells, slices were exposed to primary antibodies in blocking buffer. Secondary antibodies were diluted in 1x PBS buffer containing 0.5% BSA and 1% NDS. Samples were mounted in Prolong Gold mounting medium and coverslips were sealed with clear nail polish.

### Imaging (confocal/echo) and image analysis

Brightfield and fluorescent images of 2D cells were acquired using the Echo Revolve with the 10x objective. 2D images were analyzed using adjust Threshold (kept constant for each channel tested across images) and Analyze particles to count cells for each channel using ImageJ. Confocal microscopy for 20µm thick 3D assembled organoid slices was performed using LSM900 airyscan 2 microscope with 40X objective. Z-stacks were obtained at a step size of 1µm. Representative images were created by generating maximum intensity projections of Z-stacks using ImageJ.

### Statistics

Graphpad Prism version 10 was used to generate graphs and perform statistical tests such as One-way or two-way ANOVA with multiple comparisons tests (Holm-Sidak/ Sidak, as indicated) and Welch t-tests, as indicated. All immunostaining experiments were repeated 2-3 times independently. For each experiment, 3-4 technical replicates were used and 2-4 fields were imaged from each replicate, as indicated for individual experiments in the Figure legends.

### Quantitative RT-PCR

Quantitative RT-PCR was performed using SYBR Green detection using the SsoAdvanced Universal SYBR Green Supermix reagent (1725271) with the Bio-rad CFX96 system. Biological triplicates of cell samples (∼1-2 million cells per sample) were lysed in Trizol (Sigma, 3809). The mRNA was isolated using the Direct-zol mRNA kit (Zymo, R2050) and the cDNA was generated via the iScript kit (Bio-rad, 1708890). Primers were obtained from IDT.

### Neuron isolation and single-cell sequencing

Isolation of neurons was performed by modifying a previously published protocol (Jerber et al., 2020) as follows. Dissociation Buffer was prepared by adding Accutase (Stem Cell Technologies, 07920) to DPBS (no Calcium, no Magnesium) (Gibco, 14190144) at a ratio of 1:1 (v/v) and using 5mL of this solution to resuspend 1 vial of Papain (Worthington Biochemical Corporation, LK003176). To dissociate cells, the media was first removed, and cells were washed once with DPBS (no Calcium, no Magnesium). After aspirating, 0.5 mL of Dissociation Buffer was added to each well of the 24-well plate and incubated at 37°C in a tissue culture incubator (5% CO2) for 10-20 minutes. After 10 minutes, the dissociation process was observed periodically by gently pipetting 100 μL of Dissociation Buffer against the cells, and continuing digestion as needed. Approximately 5 minutes before the end of dissociation, Wash Buffer 1 was prepared by adding 10 μM ROCK inhibitor (Y-27632) and 1 vial DNase (D2) (Worthington Biochemical Corporation, LK003170) to 15 mL DMEM:F12. 2 mL of Wash Buffer 1 was added to each well and a P1000 pipette was used to gently agitate the cells into a single cell suspension. Cells were transferred to a 15 mL conical tube, centrifuged at 300g for 5 minutes, and then resuspended in 1 mL Cell Suspension Buffer (CSB), which consisted of 400 μg/mL BSA in DPBS (no calcium and magnesium). Cells were filtered through a 40 μm cell strainer, centrifuged at 300g for 5 minutes at room temperature, resuspended in 200 μL CSB, and gently mixed 10-15 times to ensure complete cell resuspension. Cell viability was evaluated using 0.4% Trypan Blue (Invitrogen, 2936375) and quantified with a Countess 3 automated cell counter (Invitrogen). Sample preparation and library generation were performed using the Fluent Biosciences PIPseq T2 3’ Single Cell RNA kit v4.0 PLUS. 5,000 cells per sample were targeted for sequencing. Four biological replicates were completed for each sample. cDNA fragment analysis was performed using the Agilent Bioanalyzer 2100 with the High Sensitivity DNA Kit (Agilent, 5067-4627). Libraries were pooled and sequenced using an Illumina NovaSeq X 25B 150PE with 5% PhiX at a read depth of 20,000 reads per cell (Clark et al., 2023). Sequencing data were quality controlled, aligned, and processed into count tables using PIPseeker (v3.3.0), and then loaded into Seurat (v5) for further analysis.

### PIP-seq single-cell RNA data analysis

#### Data Processing, Batch Correction, Clustering, and Annotation

Seurat R package (Version 5) was used for data processing and clustering steps. For each sample, the count matrix was filtered based on a minimum of 200 expressed genes per cell and a minimum of 3 cells with detected expression per gene. The replicates at each time point were then merged using the “merge” function of Seurat. Cells expressing a higher percentage of mitochondrial and ribosomal genes were also filtered out. We then applied sctransform normalization followed by the standard clustering workflow (RunPCA, RunUMAP, FindNeighbors, and FindClusters). We tried both the default PC number of 30 and higher (for RunUMAP and FindNeighbors) to explore the heterogeneity of the developing cell types at timepoints. We found a limited batch effect using this procedure (Figure S2). We first grouped the cells to a broad class level, such as progenitor and (induced) dopamine (DA) neurons, based on the expression of canonical progenitor cell markers, cell cycle genes, and DA neuron markers (Figure 5A, Figure S4A).

#### Per cell Midbrain score calculation

To assess the extent to which our induced neurons belong to the midbrain lineage, we compiled a list of genes as a midbrain signature and calculated a per-cell midbrain score based on the expression pattern of these genes for samples at each time point. More specifically, we used our midbrain gene list as input for “features” argument to the “AddModuleScore” function of the Seurat package (ctrl = 5, nbin = 12), which computes the average expression of the genes in the midbrain gene list for each cell, generates a control set (set of randomly selected genes from the same expression bin as the midbrain genes to normalize the module score), subtracts the background signal (mean expression of control genes) from the actual gene set expression, and returns the score. Then the final score was used to represent the relative enrichment of the midbrain gene set in each cell (Figure S4D).

#### Comparison to published midbrain and DA datasets

To investigate the correlation of our cells to in vivo midbrain and dopamine cells, as well as in vitro cells derived from published protocols, we compared the transcriptomic profiles to adult human dopamine neurons (Kamath et al., 2022) and annotated dopamine cells from cultured human midbrain organoid (Kim et al., 2024). To this end, we first compiled or re-analyzed the published datasets to obtain the cell type classification of the reference dataset. More specifically, for the adult human DA neuron dataset, we obtained the filtered count matrix and assembled the Seurat object using the “CreateSeuratObject” function, followed by integrating the downloaded metadata information (including the UMAP coordinates with the “CreateDimReducObject” function and cell type labeling for each cell). For the human midbrain organoid dataset, we obtained the barcodes.tsv.gz, features.tsv.gz, and matrix.mtx.gz files from GEO (GSM8122885), and constructed the seurat object using “Read10X” and “CreateSeuratObject” functions. Clustering was then performed using the standard workflow (‘NormalizeData’, ‘FindVariableFeatures’, ‘ScaleData’,’RunPCA’, ‘ElbowPlot’ for PC number selection, ‘FindNeighbors’ with dims of 15, ‘FindClusters’ with resolution of 1, and RunUMAP with dims of 15). To annotate this dataset, we compiled a list of marker genes for cell classes/types (Nb, DA, Nprog, Rgl, Prog, Endo/Peri and RN) as presented in the corresponding paper. Next, to compare our dataset to these reference datasets, we applied an xgboost based modified machine learning framework as described previously (Chen and Guestrin, 2016; Hahn et al., 2023; Nimkar et al., 2025; Peng et al., 2019; Wang et al., 2024; Zhang et al., 2024) to correlate the transcriptomic profile of each cell between our dataset and the reference datasets. Briefly, we subset the log-normalized expression matrices of two datasets in comparison to the shared highly variable genes (HVGs) and used 60% of the training data cells to train an XGBoost model, whose performance or accuracy was assessed on the remaining 40% of the training data cells. More specifically, a multiclass XGBoost classifier was trained using a soft-probability objective function (multi:softprob) with log-loss (mlogloss) as the evaluation metric. The model was trained for 200 boosting rounds with a learning rate (η) of 0.2, a maximum tree depth of 6, and a subsampling rate of 0.6 to reduce overfitting. The number of output classes was set to the number of unique training labels. Training categories/labels among all with highest probability value were assigned to each cell as the primary prediction. Confirming the high accuracy in the model’s prediction ability, we then use the model to predict a label for each cell in the test dataset. Finally, we summarized the results using a confusion matrix (Figure 5D, 5H).

### Multielectrode array recording

The MaxOne Chip – PSM (MaxWell Biosystems AG) are treated as per the manufacturer’s instructions. Briefly, the chips were cleaned with 1% Terg-a-zyme solution (10g/L) for 2 hours and further sterilized with 70% ethanol for 30 minutes and rinsed with deionized water after each step. For preconditioning, the chip is filled with 0.6mL culture media, covered with autoclaved MaxOne Lid and incubated in a humidity chamber in a 5% CO2 incubator at 37℃, relative humidity >95%, for 2 days, then wash it with sterile deionized water. To plate 3D organoids, the chips were coated with 0.01% Poly-L-Ornithine (PLO) solution. The next day, after rinsing the PLO and drying, 10ul of 1mg/ml Laminin is added onto the center of the chip and three 4-weeks-old 3D organoids were carefully placed above the electrodes. The 3D organoids were allowed to attach for 1-2 minutes, after which 50 μL of complete culture media (as above) is added onto the organoids. The chips with organoids were placed inside the humidity chamber into incubator for 2 hours in 5% CO2, 37℃, relative humidity >95% condition. After incubation, the well is slowly filled up to a total volume of 1 mL with complete culture media, then place it in an incubator. Culture media is replaced every 4 days. MEA recordings were performed once weekly, the day after the media change.

The chips with organoids were mounted on the MaxOne recording unit. Spike activity was recorded at a sampling rate of 20k Hz at room temperture. Band pass filtering between 300 Hz and 3kHz was applied online to reduce noise by the recording software. For the activity scan, we scanned the electrical signal from all 264000 electrodes for 30s. An online spike detector was applied to detect the spikes of each electrode. The signals that were above the 4.5 x RMS (root square mean) noise values were classified as action potentials. The activity scan is to identify the position and activity of cells on the array through sequential scanning. After activity scan, we chose 1024 channels with the highest amplitude or firing rate to record for 300s for network scanning. The network scanning was used to observe whether synchronous firing occurs. For axon tracking, we record 50 channels with the highest amplitude or firing rate for recording and scanned 9 electrodes around each chosen channel for 60s. All the spikes were sorted by SpyKing-Circus to find the single unit and each cluster of spikes was calculated to get the electrical footprint of the axon. Spike wave forms along the axon of individual cells were plotted, and the conductance velocity was calculated. The Activity Scan, Network and Axon Tracking modules were featured in MaxLab Live software.

## Resource availability

### Materials availability

This study did not generate new unique reagents.

### Data and code availability

All codes and algorithms necessary for re-analysis of the single-cell RNA-sequencing data are publicly available and can be found in other publications (Kamath et al., 2022; Kim et al., 2024). The count matrix of scRNA-Sequencing is located on Zenodo DOI 10.5281/zenodo.17489189.

This paper does not report original code.

Further information requests can be directed to Erik M. Ullian or Kriti Chaplot.

## Acknowledgment

This work was supported by Ono Pharmaceutical Co. Ltd. (E.M.U.), Weill Neurohub (E.M.U., P.M.E., I.C.C), R01EY032197 (E.M.U.), P30EY037668 (E.M.U.), R01EY030138 (X.D.) and K99EY036489 (F.W.). Dr. Yien-Ming Kuo and Suling Wang of the Ophthalmology department core are thanked for their assistance in organoid sample preparation and Figure preparation, respectively. Dr. Marius Wernig, Dr. Wayne Poon, Dr. Alex Pollen, Dr. Claire Clelland, and Dr. Mathieu Daynac are thanked for relevant discussions about the project. Dr. Bill Skarnes, Dr. Bruce Conklin and Dr. Bria Macklin are thanked for providing the iPSC lines used in this study. The Ullian and England laboratory members are thanked for their input in the project and manuscript preparation.

## Author contribution

Conception and study design: E.M.U. and K.C. Data analysis and interpretation: K.C., L.Z., and E.M.U. Manuscript writing and editing: K.C., L.Z., Z.W., F.W, I.C.C. and E.M.U. Immunofluorescence analysis, neuron, glia and organoid culturing: K.C., J.W. and M.R. ScRNA-Seq design and experiments: K.C., I.C.C., Z.W. Electrophysiology (MEA) experiments and analysis: F.W. and K.C. Bioinformatics and scRNA-seq: L.Z., I.C.C. and K.C.

---

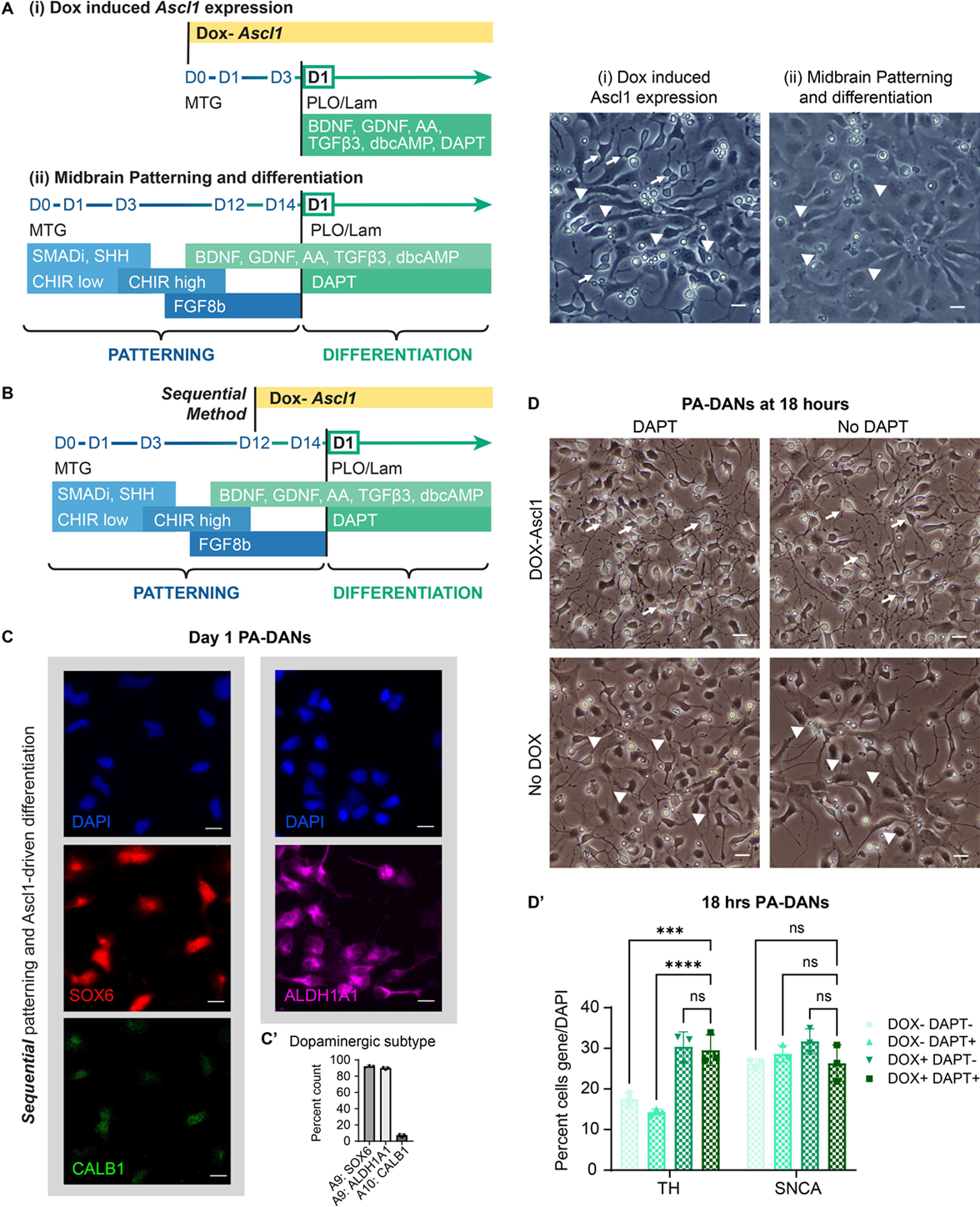

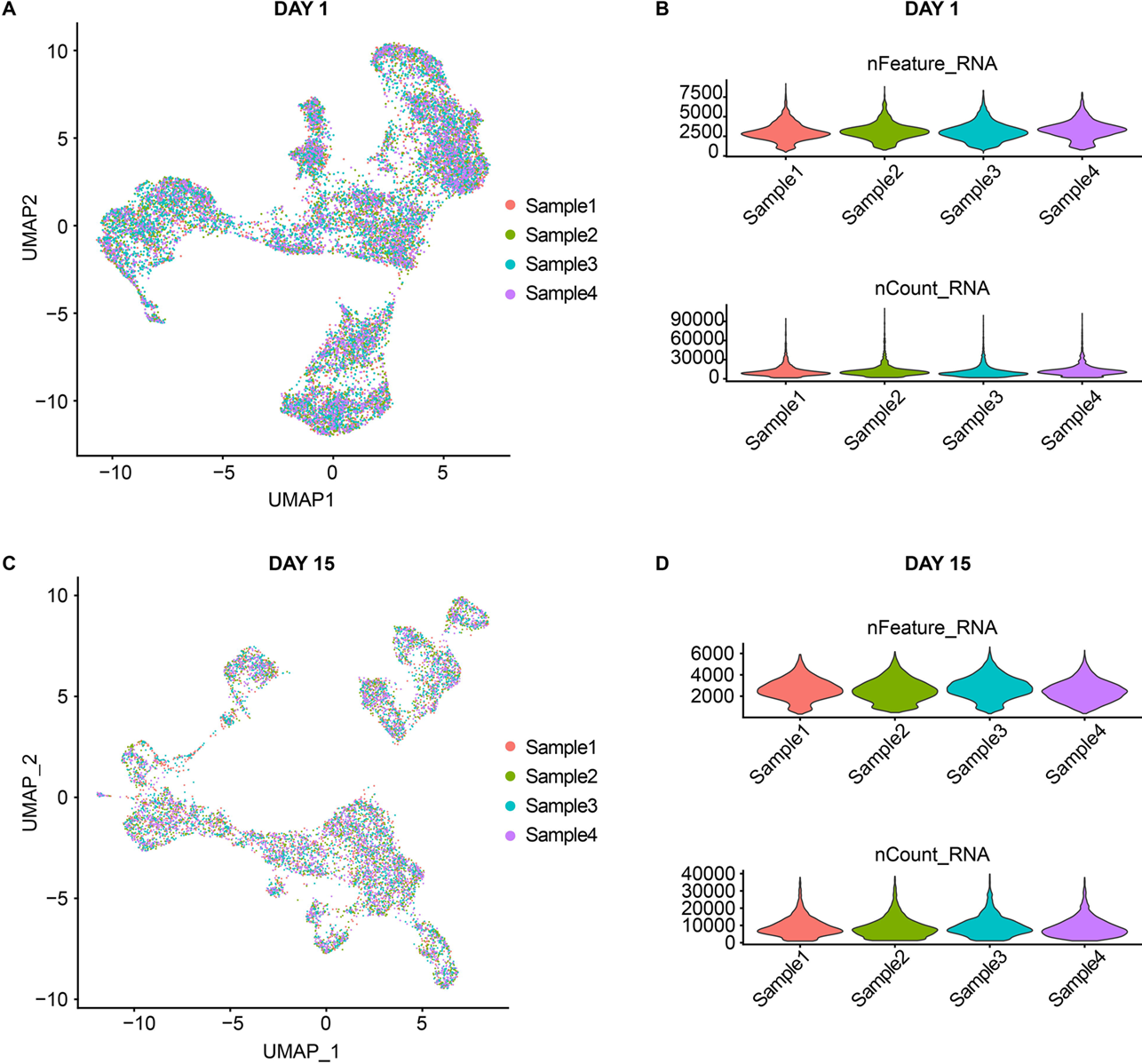

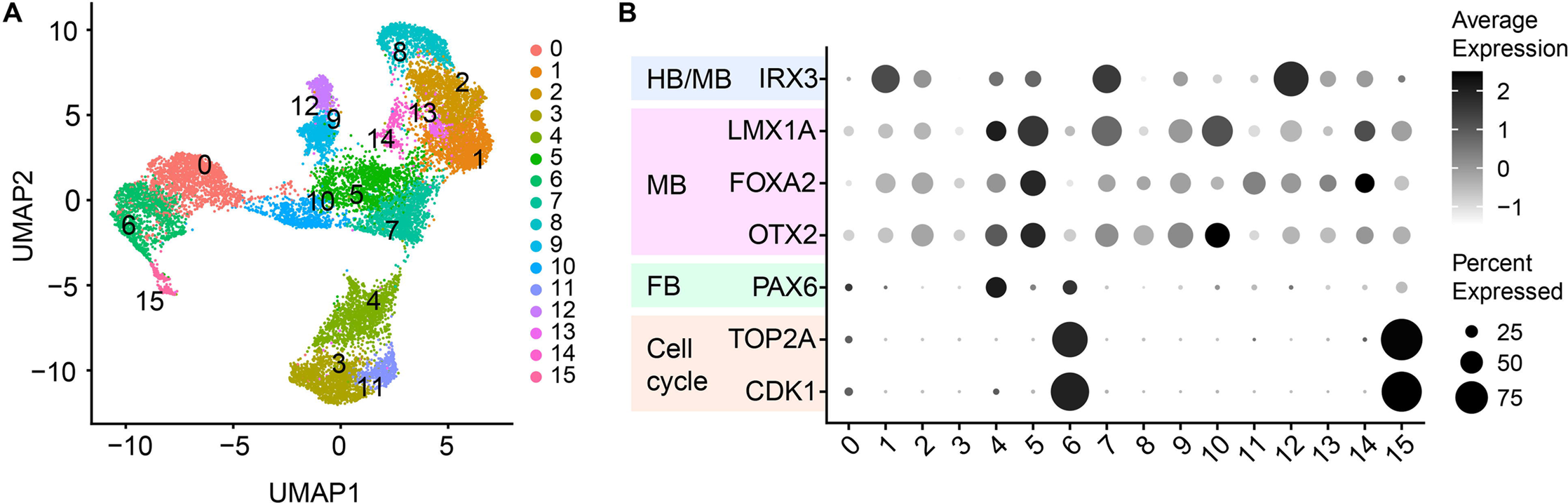

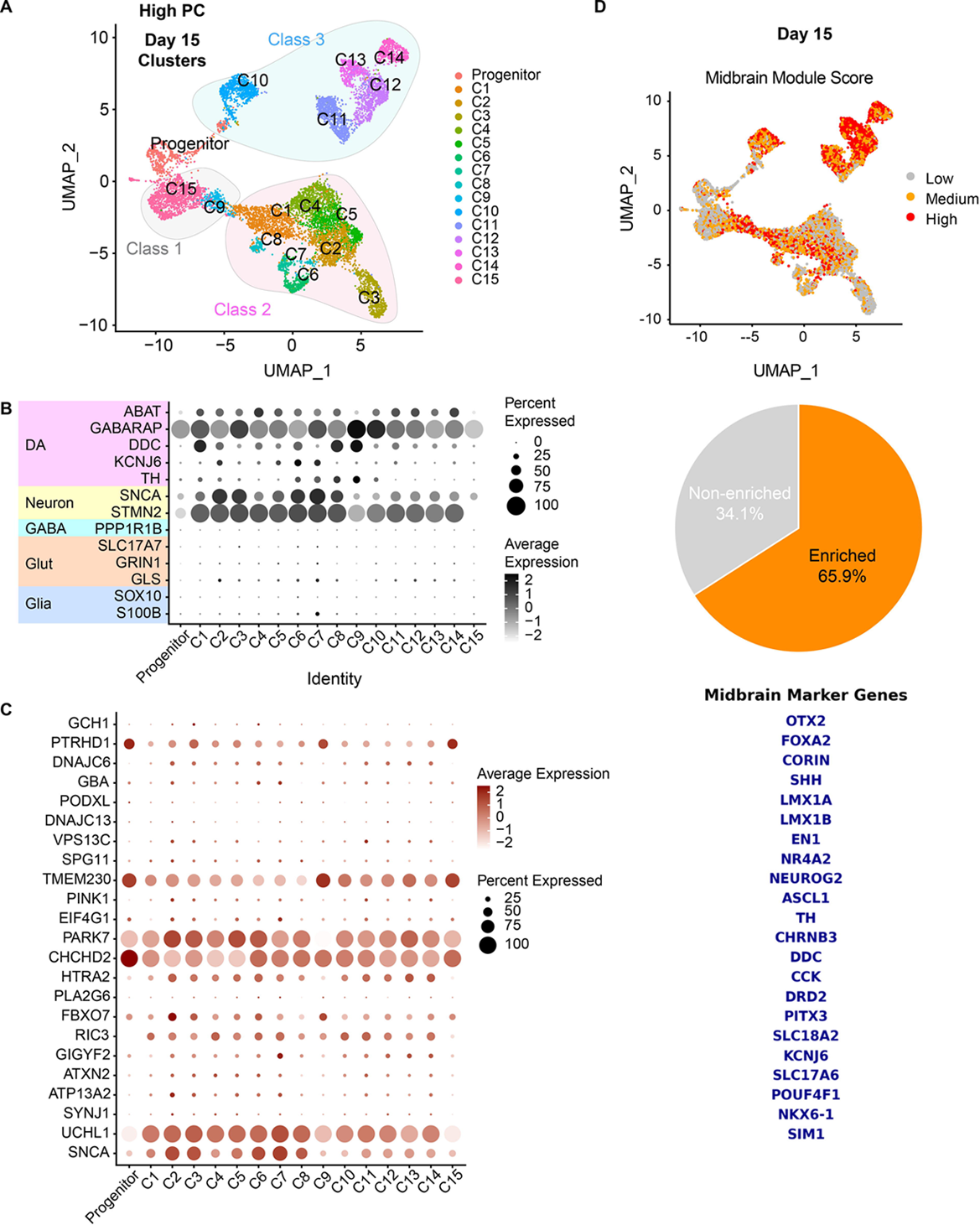

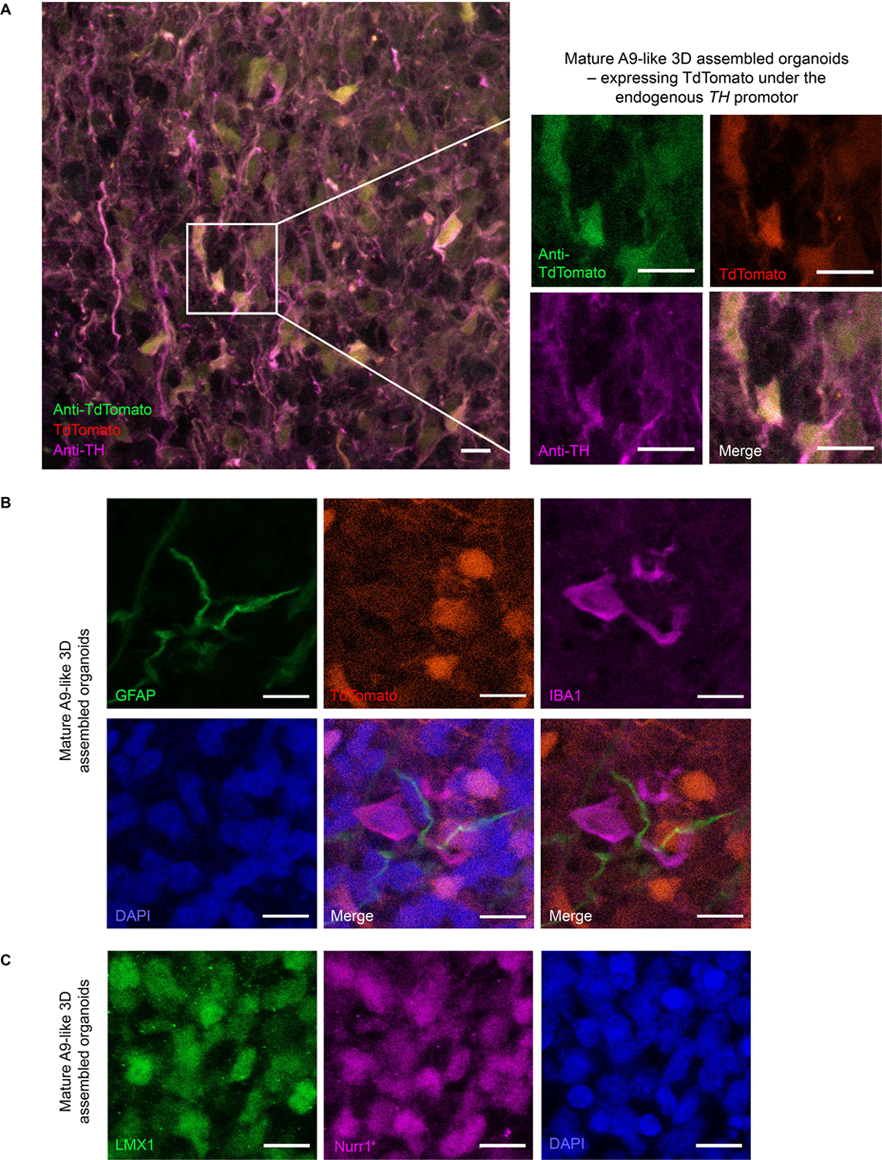

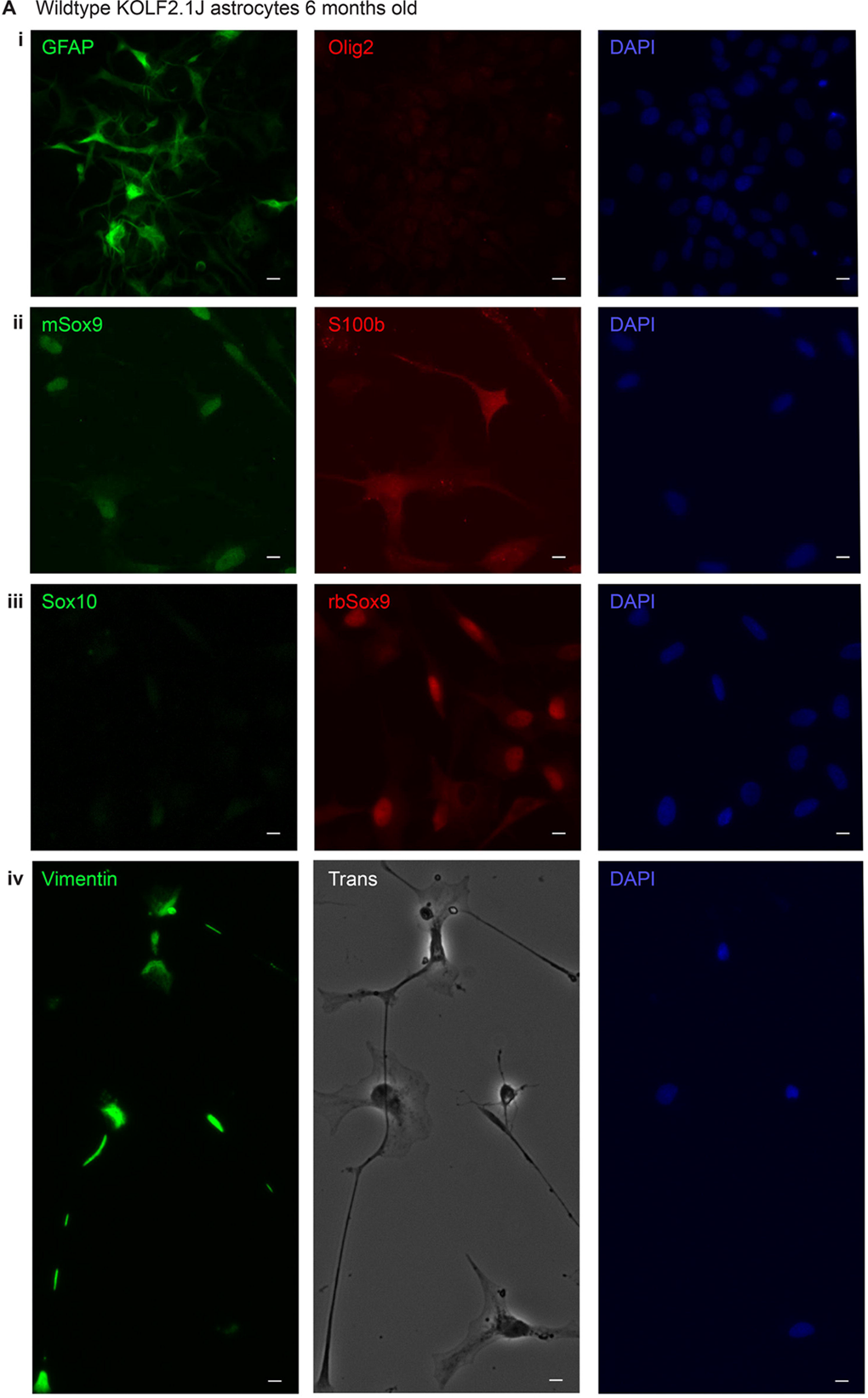

