## Supplemental information for "Rapid generation of ventral A9-like dopaminergic neurons from patterned iPSCs"

**Figure S1: Sequential patterning followed by Ascl1 expression drives DA neurogenesis and midbrain specificity.**

A. Schematic representing the timeline describing two previously established approaches (left) to generate DA neurons: (i) induction of Ascl1 in iPSCs, and (ii) directed differentiation of iPSCs using small molecules into midbrain lineage and dopaminergic neurons. Representative brightfield images (right) of cells generated via the 2 approaches after 24 hours of plating (i) Dox-induced Ascl1 expression yielding low neurogenesis, and (ii) directed midbrain-specific differentiation yielding progenitor-like cells. Arrows indicate neuron-like morphology, arrowheads indicate progenitor-like morphology. Scale bar 20μm.

B. Schematic showing a timeline for sequential method to generate patterned Ascl1-driven dopaminergic neurons (PA-DANs).

C. Representative images of PA-DANs, after 24 hours of plating, immunostained with A9-specific markers such as SOX6 (red) and ALDH1A1 (magenta) and A10 marker CALB1 (green). Corresponding nuclear staining with DAPI is shown (blue). Scale bar 10μm. Graph (C') showing high expression of A9 markers, SOX6 and ALDH1A1 and low expression of A10 marker CALB1. Error bars indicate std. dev. n=3, 3 fields from each replicate. Approx 2000 total cells counted per condition.

D. Representative brightfield images of PA-DANs, after 18 hours of differentiation, showing neuronal morphology in the presence of DOX, with (i) or without (ii) DAPT. PA-DANs differentiated in the absence of DOX, with (iii) or without (iv) DAPT, showed more non-neuronal, progenitor-like morphology. Scale bar 20μm. Graph (D') showing percentage of PA-DANs expressing TH and SNCA. TH is expressed in higher percentage of PA-DANs in presence of DOX inducer. Error bars indicate std. dev. n=3, 3 fields from each replicate. Approx 1000 total cells counted per condition.

**Figure S2. QC and filtering steps for ScRNA Day 1 and Day 15 datasets**

A. UMAP representation showing Day 1 replicates

B. Violin plot features/counts for Day 1 replicates

C. UMAP representation showing Day 15 replicates

D. Violin plot features/counts for Day 15 replicates

**Figure S3. Marker gene expression panel analysis on Day 1 shows midbrain specificity**

A. UMAP representation of Day 1 PA-DANs with 15 clusters

B. Dot plot showing expression of forebrain (FB), midbrain (MB), hindbrain (HB), cell cycle genes on Day 1 in each cluster (absence of Astrocytic and oligodendritic markers and hindbrain marker, GBX2)

**Figure S4. Transcriptomic analysis of Day 15 PA-DANs shows persistent midbrain specificity and DA neuronal differentiation**

A. UMAP representation of Day 15 PA-DANs showing clusters depicting CDK1-positive progenitors (orange) and neuronal cell types designated as Class 1, Class 2 and Class 3. Equivalence to Fig5A: Progenitors and Class 1 clusters are equivalent to Cluster 2; Class 2 is equivalent to Clusters 0 and 4; Class 3 is equivalent to Clusters 1 and 3 in Fig 5A.

B. Dot blot showing the high DA and neuronal markers and low expression of astrocytic, oligodendrocytic, glutamatergic, and GABAergic markers.

C. Dot blot showing the expression of key PD-associated genes

D. UMAP representation (top) of Day 15 PA-DANs highlighting the high, medium and low midbrain marker score, and pie-chart (middle) representing the enriched and non-enriched percentages of Day 15 PA-DANs showing high and medium midbrain module score, calculated based on the list of 22 midbrain and DA-specific genes, compared against a set of control genes.

**Figure S5: A9-like 3D organoids generated with PA-DANs containing a Td-Tomato reporter expressed under the *TH* promotor.**

A. Representative confocal image of 4 weeks old 3D organoids showing costaining using Anti-Tdtomato (green) and Anti-TH (magenta) antibodies overlapping with Tdtomato (red) fluorescence. Inset showing 2x magnified field of PA-DANs. Scale bar 10µm

B. Representative confocal images of 3 weeks old 3D organoids immunostained for GFAP (green) and IBA1 (magenta) showing mature morphology of iAstrocytes and iMicroglia. PA-DANs labelled with endogenous Td-Tomato (red) expressed under the TH promoter (generated from KOLF2.1J Line 2), are juxtaposed with GFAP-stained iAstrocytes and IBA1-staing iMicroglia. Scale bar 10µm

C. Representative confocal images of 5 weeks old 3D organoids immunostained for LMX1A/B (green) and Nurr1 (magenta) showing nuclear co-localization with DAPI (blue). Scale bar 10µm

**Figure S6: iPSC-derived astrocytes matured under 2D conditions show astrocytic markers**

A. Representative images of iAstrocytes differentiated from 6 months old KOLF2.1J astrocyte precursor cells show expression of GFAP (i, green), SOX9 (ii, green; iii, red), S100β (ii, red) and vimentin (iv, green) and lack oligodendrocytic markers Olig2 (i, red) and SOX10 (iii, green). Trans image (iv) shows expected astrocytic cell morphology and processes. Corresponding DAPI co-stain marks the nucleus. Scale bar 10µm
